# NovoLign: metaproteomics by sequence alignment

**DOI:** 10.1101/2024.04.04.588008

**Authors:** Hugo B.C. Kleikamp, Ramon van der Zwaan, Ramon van Valderen, Jitske M. van Ede, Mario Pronk, Pim Schaasberg, Maximilienne T. Allaart, Mark C.M. van Loosdrecht, Martin Pabst

## Abstract

Tremendous advances in mass spectrometric and bioinformatic approaches have expanded proteomics into the field of microbial ecology. The commonly used spectral annotation method for metaproteomics data relies on database searching, which requires sample-specific databases obtained from whole metagenome sequencing experiments. However, creating these databases is complex, time-consuming, and prone to errors, potentially biasing experimental outcomes and conclusions. This asks for alternative approaches that can provide rapid and orthogonal insights into metaproteomics data. Here we present NovoLign, a *de novo* metaproteomics pipeline that performs sequence alignment of *de novo* sequences from complete metaproteomics experiments. The pipeline enables rapid taxonomic profiling of complex communities and evaluates the taxonomic coverage of metaproteomics outcomes obtained from database searches. Furthermore, the NovoLign pipeline supports the creation of reference sequence databases for database searching to ensure comprehensive coverage. The NovoLign pipeline is publicly available via: https://github.com/hbckleikamp/NovoLign.

## INTRODUCTION

Microorganisms form complex multi-species communities in nature, inhabiting virtually every ecological niche on Earth. They are involved in global biogeochemical cycles and significantly affect human health and well-being^1-4^. Therefore, understanding these complex microbial ecosystems is required to accelerate the development of effective biotechnological solutions to address global challenges^5-8^. However, the majority of microorganisms cannot be cultured in laboratory settings, and thus require advanced culture-independent methods to study them^9-11^. Among these techniques, metaproteomics is particularly powerful as it allows to determine expressed metabolic functions, the protein biomass composition, and species-to-species interactions^11-13^. However, compared to traditional single organism proteomics, metaproteomics faces the challenge of analyzing highly complex systems with potentially hundreds of species present at varying abundances^12, 14^. Consequently, metaproteomic experiments depend on reference sequence databases that accurately cover all organisms present in the microbial community^15-19^. Current strategies for constructing metaproteomic databases involve comprehensive generic reference databases such as the NCBI nr, RefSeq or UniRef. However, large databases require focusing of the sequence space to allow sensitive metaproteomics experiments. This can be achieved by using empirical knowledge about the microbial niche or by incorporating taxonomic information from other techniques such as 16S rRNA sequencing^20-27^. More recent focusing approaches employ also multi-round searches and *de novo* sequence information^21, 25, 28-30^. Nonetheless, generic reference sequence databases may not encompass all organisms, strains, and sequence variants present in the community, which compromises accuracy and coverage. Therefore, for most applications the preferred method to create reference sequence databases is whole metagenome sequencing of the target community^12, 15, 31^. However, whole metagenome sequencing experiments are complex, expensive, and time-consuming. The coverage of the metagenomic database can be biased by the employed DNA extraction method, sequencing errors, and variations between data processing pipelines^14, 16^. Consequently, metaproteomic experiments are highly influenced by the database construction procedure. Unfortunately, there are currently no dedicated pipelines which independently evaluate the outcome of metaproteomic experiments.

However, peptide *de novo* sequencing allows to annotate mass spectrometric fragmentation spectra with amino acid sequences in a database-independent manner^32-35^. Therefore, *de novo* sequencing has been already employed where reference sequence database are not available, such as for determining the amino acid sequence of antibodies^36, 37^. *De novo* sequencing has been also used to access the suitability of reference sequence files in proteomics experiments^38^, and to string-search large public reference sequence databases to obtain taxonomic information from a microbial community^30, 39^. For example, the recently developed NovoBridge pipeline automates quality evaluation and matching of *de novo* sequences to a precomputed peptide database (Unipept) to provide rapid taxonomic information from complex samples^39, 40^. However, the quality of mass spectrometric sequencing data, *de novo* sequencing errors, and incomplete reference sequence databases limit the number of sequencing spectra that provide confident taxonomic information. Furthermore, using precomputed peptide databases require continuous update of the sequences and hamper the use of alternative proteolytic enzymes or the search for amino acid modifications.

To overcome the limitations posed by *de novo* sequencing errors and incomplete reference sequence databases, *de novo* sequence tags can also be matched to reference databases using sequence alignment. This has already been employed in tools such as PeptideSearch, CIDentify, MS-BLAST, MS-HOMOLOGY, FASTS, MS-Spider, and OpenSea, amongst others^41-48^. However, performing sequence alignment of large volumes of data against very large reference sequence databases is computationally highly demanding, and not suitable for processing complete metaproteomics experiments. In 2015, Buchfink et al. introduced the sequence aligner DIAMOND, which allows to align large volumes of data on standard desktops and eases the use of custom reference sequence databases^49^. This laid the foundation for creating a sequence alignment tool capable of handling large volumes of metaproteomic sequencing data with custom databases.

In this study, we introduce NovoLign, a *de novo* metaproteomics pipeline that classifies de novo sequences from complete metaproteomic experiments using sequence alignment against very large reference sequence databases. The pipeline aligns *de novo* sequences (including decoys) from complete metaproteomics experiments against generic reference sequence databases in short time frames on standard desktop PCs. The sequence alignment pipeline overcomes challenges posed by *de novo* sequencing errors and incomplete reference sequence databases. The fraction of false positive classifications are estimated using randomized sequences. This metagenomics-independent approach allows to perform rapid taxonomic profiling with deep coverage, and to evaluate the sequence coverage of conventional metaproteomics experiments that employ database searching. Furthermore, the obtained taxonomic composition can be used to construct or complement reference sequence databases. We demonstrate the performance of NovoLign with a large spectrum of pure reference strains, enrichment cultures, synthetic and complex natural communities.

## MATERIALS AND METHODS

### Application of publicly available data

Employed pure reference strains, enrichment cultures synthetic and natural communities, taxonomic lineages and content of synthetic community samples are summarized in SI Table 3. Briefly, the equal protein synthetic community proteomic raw data and reference database were obtained from ProteomXchange server project PXD006118 (Kleiner et al., 2017, Nat Commun)^50^, the simplified human gut microbiota model (SIHUMIx) proteomic raw data and reference database were obtained from ProteomXchange server project PXD023217 (Van den Bossche et al., 2021, Nat Commum)^12^. *Acinetobacter baumannii* raw data and reference database were obtained from PXD011302 (Di Venanzio et al., 2019, Nat Commun)^51^, *Caldalkalibacillus thermarum* from PXD042369 (de Jong et al., 2023, Front Microbiol)^52^, *Nitrospira moscoviensis* from PXD019583 (C. Lawson et al., 2021, mSystems)^53^, *Chlamydomonas reinhardii* from PXD010160 (Scholz et al., 2019, Plant Journal)^54^, *Lactobacillus sakei* from PXD011417 (Prechtl et al., 2018, Front Microbiol)^55^, *Paracoccus denitrificans* from PXD013274 (Schada von Borzyskowski, 2019, Nature)^56^, *Streptococcus mutans* from PXD006735 (Ahn et al., 2017, Scientific Reports)^57^, *Halanaeroarchaeum* sp. HSR-CO from PXD028241 (Sorokin et al., 2022, The ISME Journal)^58^, the *Clostridium kluyveri*-dominated enrichment, preparation and analysis was performed as described in Allaart et al., (2021), Front. Bioeng. Biotechnol.^59^, and Allaart et al., (2023) Scientific Reports PXD040972^60^. The raw data for the following samples are provided via PXD050548. More specifically, the *Saccharomyces cerevisiae* tryptic digest was purchased from Promega (Cat No. V7461) and analyzed as described in Pabst et al., The ISME Journal, 2020^61^, except using a shorter one-dimensional 60-minute gradient for chromatographic separation. The preparation and analysis of *Aeromonas bestiarum* is described in Tugui et al., 2024, bioRxiv^62^. The *Ca*. Kuenenia stuttgartiensis enrichment was prepared and analyzed as described in Lawson et al., 2019, The ISME Journal^63^. The *Ca*. Accumulibacter phosphatis enrichment was prepared and analyzed as described in Kleikamp et al., 2021, Cell Systems^39^. Aerobic granular sludge was sampled from the wastewater treatment plant in Utrecht, The Netherlands, and prepared and analysed as described in Kleikamp et al., Water Research, 2023^14^. All analysed proteomics and metaproteomics reference sample data are summarized in SI EXCEL Table 1.

### *De novo* sequencing and database searching of mass spectrometric raw files

The mass spectrometric raw data were processed using PEAKS Studio X (Bioinformatics Solutions Inc., Canada)^64^ for database search and *de novo* sequencing, or DeepNovo^65^ for obtaining additional *de novo* sequencing output files for developing the pipeline. Both, *de novo* sequencing and database searching was performed allowing 20 ppm parent ion and 0.02 Da fragment mass error. Carbamidomethylation was set as fixed and methionine oxidation as variable modifications. Database searching was performed with N/Q deamidation as additional variable modifications. Database searching further used decoy fusion for estimation of false discovery rates (FDR) and subsequent filtering of peptide spectrum matches for 1% FDR. Only the top ranked *de novo* sequence annotations were considered for processing. Proteome reference sequence databases for comparative database searching were obtained from the provided ProteomeXchange server projects or published supplementary data. The proteome reference database for the *Clostridium kluyveri* dominated enrichment and the wastewater sludge was obtained from whole metagenome sequencing experiments. The aerobic granular sludge was fractionated by size and then homogenized, following the protocol described in Kleikamp et al., 2023, Water Research^14^. For both samples (*Clostridium kluyveri* enrichment and wastewater granule fractions), DNA was then extracted using a DNeasy UltraClean Microbial Kit (Qiagen, Germany), and the extracted DNA was quantified with a Qubit fluorometer. Whole metagenome sequencing was performed on an Illumina NovaSeq platform with paired-end reads (Novogene Co. Ltd., China). Raw reads were quality checked, trimmed and then assembled using MEGAHIT (v1.0.4-beta). Thereafter, ORFs were predicted with MetaGeneMark (v3.05) using default parameters, for scaftigs ≥500 bp. Finally, redundancy in the predicted ORFs was eliminated using CD-HIT (v4.5.8). The individual metagenomic databases of the granule size fractions were merged before employing a two-round database search approach. All other samples, including the *Clostridium kluyveri* enrichment, were analysed using the above outlined database searching procedure.

### Training sequences with *de novo* errors and mutations

The developed Python code and additional documentation to generate peptides with simulated *de novo* sequencing errors and mutations are freely available via GitHub: https://github.com/hbckleikamp/De-Novo-ErrorSIM/. *De novo* error files were created for “equal mass substitutions”, “inversions of amino acids”, and “Other” mutations^33^. Equal mass substitutions involve substituting a combination of one or more amino acids with another combination of amino acids that have the same mass. This, for example, includes the substitution of asparagine (N) with two glycine residues (GG). Equal mass combinations of up to 6 amino acids were created, where a sliding window detected substitutable regions within a peptide, which are then selected for substitution at a 25, 50 and 100% chance. For inversions of 2 or 3 amino acids, the amino acids remains same, but their sequence order was scrambled at a 1, 5, 10 and 20% chance. The “other” peptides included mutated peptides, which were generated by reverse translation into codons, followed by random base mutation at 1, 5, 10 and 20% chance. Thereby, larger peptides are more likely to contain multiple *de novo* sequencing errors. To make sure that each error type generated similarly different peptides, the Levenshtein distance between the original and altered peptide was computed^66^. While inversions and mutations can occur in any position, equal mass substitutions can only occur if certain combinations of amino acids are present. Therefore, if similarly “different” peptides need to be generated, they need to occur at a higher error rate. Various error rates were tested for the different error types to create a similar distribution to 5% mutation rate. Final error rate was selected as 5% for mutation and inversion, and 25% for substitution errors. The individual *de novo* error sequences were combined using rates of occurrence for the individual error types as observed for common sequencing tools recently^33^ (substitution of 1 by 1 or 2 AAs = 6.3%, substitution of 2 by 2 AAs = 13.7%, substitution of 3 by 3 AAs = 6.3%, substitution of 2 by 3 AAs = 3.7%, substitution of 4 by 4 AAs = 9.7%, substitution of 5 by 5 AAs = 7.6%, substitution of 6 by 6 AAs = 6.8%, inversion of 2 or 3 AAs = 16.1%, other = 29.7%).

### Outline of NovoLign pipeline

The NovoLign pipeline is a single “tunable” python script in which parameters can be altered manually. The pipeline can be used with the commonly employed proteomic reference sequence databases including NCBI, UniprotKB and GTDB. The pipeline, additional documentation and example data are freely available via GitHub: https://github.com/hbckleikamp/NovoLign. Setup of the pipeline and databases is described in the online documentation of the GitHub page. NovoLign was tested with output file formats from PEAKS Studio X^64^ and DeepNovo^65^. Nevertheless, any tabular or .txt-like format can be supplied, provided it contains a column of peptide sequences with the header “Peptide”. Supplying database-searched peptides (PSM file) and a reference sequence database (FASTA file with NCBI TaxIDs) will furthermore allow the evaluation of spectral and database coverage. The pipeline comprises the following modules:

1. DIAMOND alignment (write_to_fasta.py, diamond_alignment.py): Firstly, peptide sequence lists are imported into the Python environment and filtered based on *de novo* score thresholds. Default filters include a minimum quality score of 70 ALC% for PEAKS data or −0.1 minimum score for DeepNovo. Optional (additional) filters include maximum ppm mass error, minimum peptide length, and peak intensity or area. Filtered sequences are written to a fasta file. Decoy peptide sequences are created by scrambling the order of amino acids in front of the cleavage site (R or K) of every sequence. The resulting peptide file is aligned using the sequence aligner DIAMOND^49^, using parameters optimized for *de novo* sequencing errors. Default parameters include: a minimum percent identity of 85%, a minimum coverage of 80%, and a PAM70 matrix with gap opening penalty 2 and gap extension penalty of 4.
2. Lowest Common Ancestor analysis (process_alignment.py): Output files from sequence alignment are filtered for a minimum bitscore of 25. The obtained taxonomy IDs are then mapped to lineages for subsequent lowest common ancestor (LCA) analysis. NovoLign includes 3 algorithms for LCA analysis, which contain different levels of stringency and false positives. The conventional LCA algorithm (CON) has the highest stringency and retains all aligned sequences. The weighted LCA algorithm (W)^67^ counts the frequencies of all aligned taxonomies, which act as weights. For each peptide, the taxonomic weights of aligned sequences are sorted, cumulatively summed, and normalized to 1. The most frequent taxonomies are kept for LCA until the “weight_cutoff”(default 0.6) is reached. The bitscore LCA (BIT), operates similarly, but instead groups the aligned sequences by their taxonomic lineages for each ranks and sums the bitscores. Only taxa with a weight over the “weight_cutoff”(default 0.6) are retained^39, 68^.
3. Taxonomic composition (bar_graphs.py): Once the LCA has been determined, a frequency cutoff “freq_cut” (default 5) is applied, in order to remove taxonomies that occur at very low frequency. The remaining taxonomies are grouped together to provide a taxonomic composition of the analyzed microbial community. The results are then written to a table and displayed in bar graphs. Furthermore, the sequences from the target database (against which the query peptides were aligned) can be collected to create a specialized protein reference database. Optional arguments include an exclusion list, addition of decoy sequences, or addition of the query *de novo* peptides to the database. Alternatively, all sequences that belong to the identified taxonomies (at different levels e.g. species, genus, family) can be extracted and compiled into a database.
4. Spectral quality analysis (experiment_qc.py): For spectral quality control, aligned *de novo* peptides are visualized with a scatter plot, histograms, and a stacked bar chart, based on *de novo* scores and annotation rates. Aligned *de novo* peptides are grouped into: “exact”, (100% identity, 100% coverage), “exact tag” (100% identity, <100% coverage), “aligned” (< 100% identity, 100% coverage), “aligned tag” (<100% identity, <100% coverage) and unmatched spectra. Optionally, if database searched peptides are also supplied, the stacked bar chart will show if the spectra of *de novo* sequenced peptides are detected in database searching (“matched” or “not detected”).
5. Analysis of reference sequence database (database_qc.py): To access the coverage of the reference sequence database used for database searching, the aligned *de novo* sequences are compared to the database searching outputs. The annotated taxonomies will be compared for the ranks: order, family, genus and visualized with stacked bar charts. Taxonomic distributions will be shown for peptide spectrum matches detected exclusively with *de novo* sequencing (DN_only), all *de novo* peptides (DN_all), peptides detected exclusively in database searching (DB_only) and all database searching peptides (DB_all).

## RESULTS

Here we present NovoLign, a *de novo* metaproteomics pipeline that performs large-scale sequence alignment of *de novo* sequences from complete metaproteomics experiments against large reference sequence databases. To optimize the alignment and post-processing parameters, as well as to evaluate the accuracy of the taxonomic profiling, we employed a wide range of in silico and experimental data. These included *de novo* error datasets, pure reference strains, laboratory enrichments, and samples from synthetic and natural microbial communities.

### Sequence alignment of *de novo* sequences

The sequence alignment of *de novo* sequences presents two major challenges. Firstly, bottom up proteomics experiments result in short peptide sequences which are only poorly aligned using DIAMOND default settings. Secondly, in practice, many mass spectrometric fragmentation spectra show gaps in the fragment ion series and misleading fragments, resulting in sequencing errors such as amino acid inversions or substitutions^33, 35^. However, many of these errors occur in blocks within the amino acid sequence, and thus may still provide alignment to the correct taxonomy. To evaluate the correct alignment of such short sequences and sequences with errors, we created 13 in silico datasets, with 1000 sequences each (SI DOC A and SI DOC Table 1), to reflect commonly observed sequencing errors like different types of amino acid inversions, substitutions, and mutations. Additionally, we constructed a dataset that combined all these errors, with error rates as commonly observed for the individual error types in proteomics experiments^33^ (SI DOC Figures 1 and 2). This combined error dataset was then used to determine the most suitable DIAMOND parameters. First, we compared the alignment performance when using different scoring matrices, seed shapes, and DIAMOND seed search algorithms. Generally, for short sequences, PAM substitution matrices are more suitable than BLOSUM matrices (SI DOC, section B). Shevchenko et al. (2001) used a PAM30 scoring matrix in their MS BLAST tool^43^. To improve the reporting of highly similar sequences, the authors made modifications to the matrix, such as adding scores for isobaric amino acids, trypsin cleavage sites, and substitutions. However, a modified matrix may interfere with the identification of homologous sequences from related organisms, which could be a disadvantage for metaproteomics experiments where the analyzed organisms may not be present in the database. Therefore, we accounted in our pipeline only for the isobaric amino acids isoleucine (I) and leucine (L) by replacing all “I” with an “L” in our reference sequence databases and sequencing data, but we did not modify scoring matrices. Additionally, the DIAMOND default sensitivity mode employs seeds with lengths of 12 and 15, which may discriminate shorter sequences. Furthermore, the different seed search algorithms (i.e. 0 = double-indexed, default algorithm, for large input files, less efficient for small query files; 1 = query-indexed, better performance for small query files; ctg = contiguous-seed mode, further improved performance for small query files) can affect the alignment of short sequences (with errors) and smaller datasets^69^. When aligning the combined error dataset against the UniRef100 and Swiss-Prot databases, we found that (as expected) the PAM 30 and 70 matrices generally performed better than BLOSUM62. Additionally, using shorter custom seeds and the “ctg” mode improved alignment for shorter peptides (<10 amino acids) significantly (SI DOC Figures 3 and 4). Furthermore, standard scoring matrices often use high gap opening and low gap extension penalties. This is suboptimal when aligning *de novo* peptide sequences with errors. Therefore, we evaluated different combinations of gap opening and gap extension penalties and found that a 2/4 (gap opening/extension penalties) combination performed best for our established in silico *de novo* sequences (SI DOC Figures 5 and 6). Finally, when comparing different reporting parameters for “query cover” and “percentage of sequence identity”, we found that a combination of % identity 85 with a % query coverage in the range of 75–85 provided the highest accuracy (SI DOC Figures 7 and 8). After determining the most suitable DIAMOND parameters we investigated the alignments when working with larger reference sequence databases, such as UniRef100. The main objective was to determine the fraction of sequences which provide i) exact sequence matches, ii) correct taxonomy matches, iii) false-positive matches, iv) random matches, and v) unmatched sequences, as well as to investigate the length distribution of the aligned sequences. Remarkably, the combined error dataset provided for a large fraction of sequences exact or correct matches (SI DOC Figures 9–12). For example, at the lower ranks order, family and genus, we obtained approx. 60% alignments to the correct taxonomy for Swiss-Prot and approx. 50% for UniRef100. However, most importantly, the fraction of sequences that provided alignments to wrong taxonomies was <10%. Furthermore, the length distribution of the aligned sequences reflected the distribution of the input sequences, with only a minor discrepancy towards smaller peptides with <10 amino acids. The UniRef100 database provided more alignments for the random (decoy) sequences. These, however, generally show lower bitscores and can thus be eliminated by setting a minimum bitscore threshold. Overall, the individual *de novo* error datasets provided outcomes comparable to the combined dataset (SI DOC Figures 13–23). However, the randomly mutated sequences (datasets 12 and 14, as shown in SI DOC Table 1 and SI DOC Figure 14) yielded significantly fewer alignments. Additionally, sequences that underwent substitutions involving three or more amino acids (datasets 31–42, detailed in SI DOC Table 1 and SI DOC Figures 20–23) exhibited increasing discrimination against shorter sequences. In summary, the optimized DIAMOND parameters allowed to align and retrieve correct taxonomies for a large fraction of *de novo* peptides with common sequencing errors and different lengths (10–50 amino acids). Most importantly, erroneous annotations were <10%, which makes these alignments suitable for metaproteomic applications.

**Figure 1.**
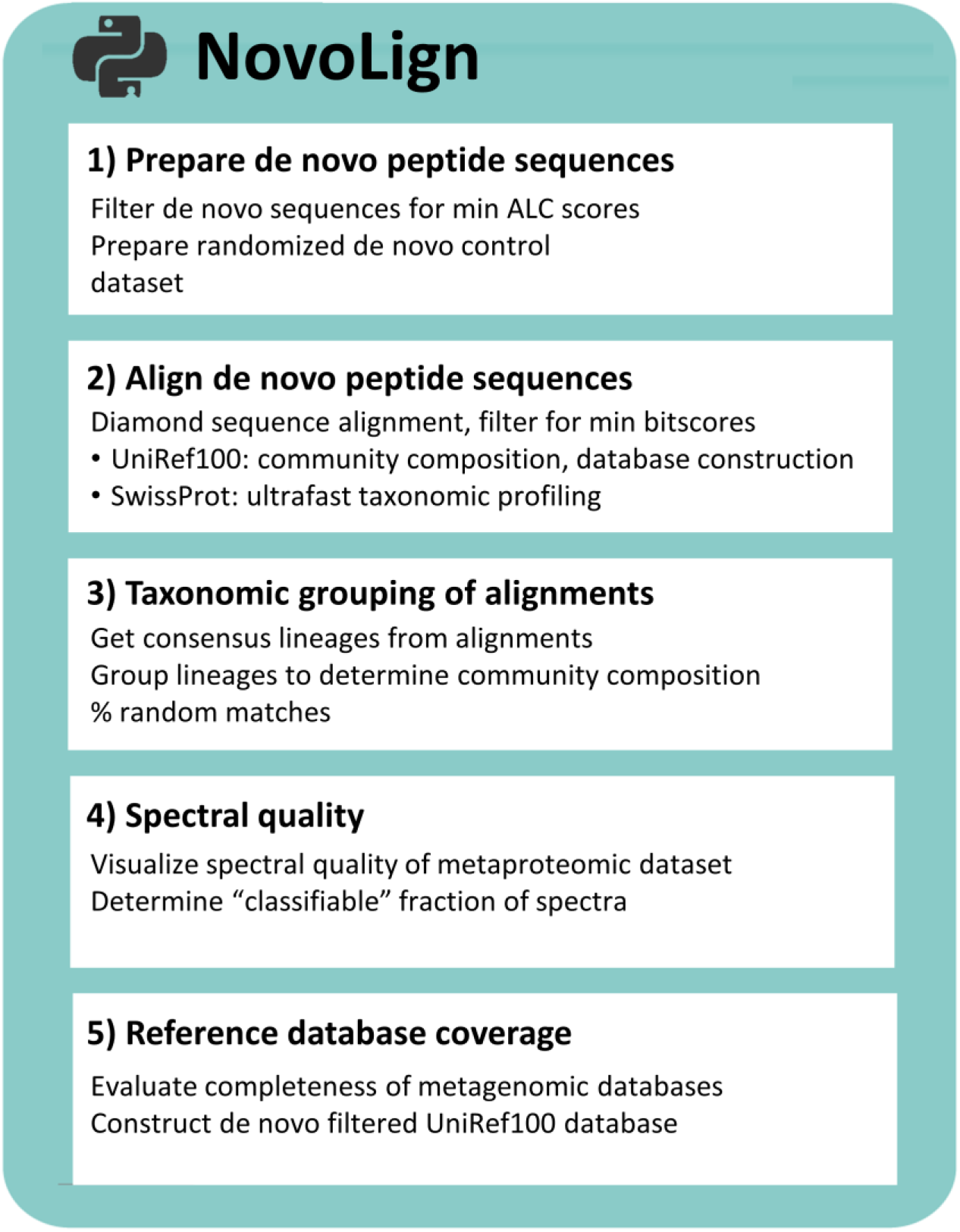
The table outlines the main modules of the NovoLign pipeline, which conducts large-scale sequence alignment of complete metaproteomics experiments. This enables rapid (database searching independent) taxonomic profiling with deep coverage. It also offers an alternative approach for assessing the quality and coverage of metaproteomics outcomes obtained from database searching approaches. The pipeline contains 5 modules: 1) preparation of *de novo* sequences and generation of randomized decoy sequence dataset, 2) alignment of the *de novo* and decoy sequences using DIAMOND, 3) taxonomic grouping of the aligned sequences and determining the percentage of random matches, 4) evaluating the spectral quality of the analyzed metaproteomics experiment, and 5) evaluating the coverage of the reference sequence database used for database searching.

**Figure 2.**
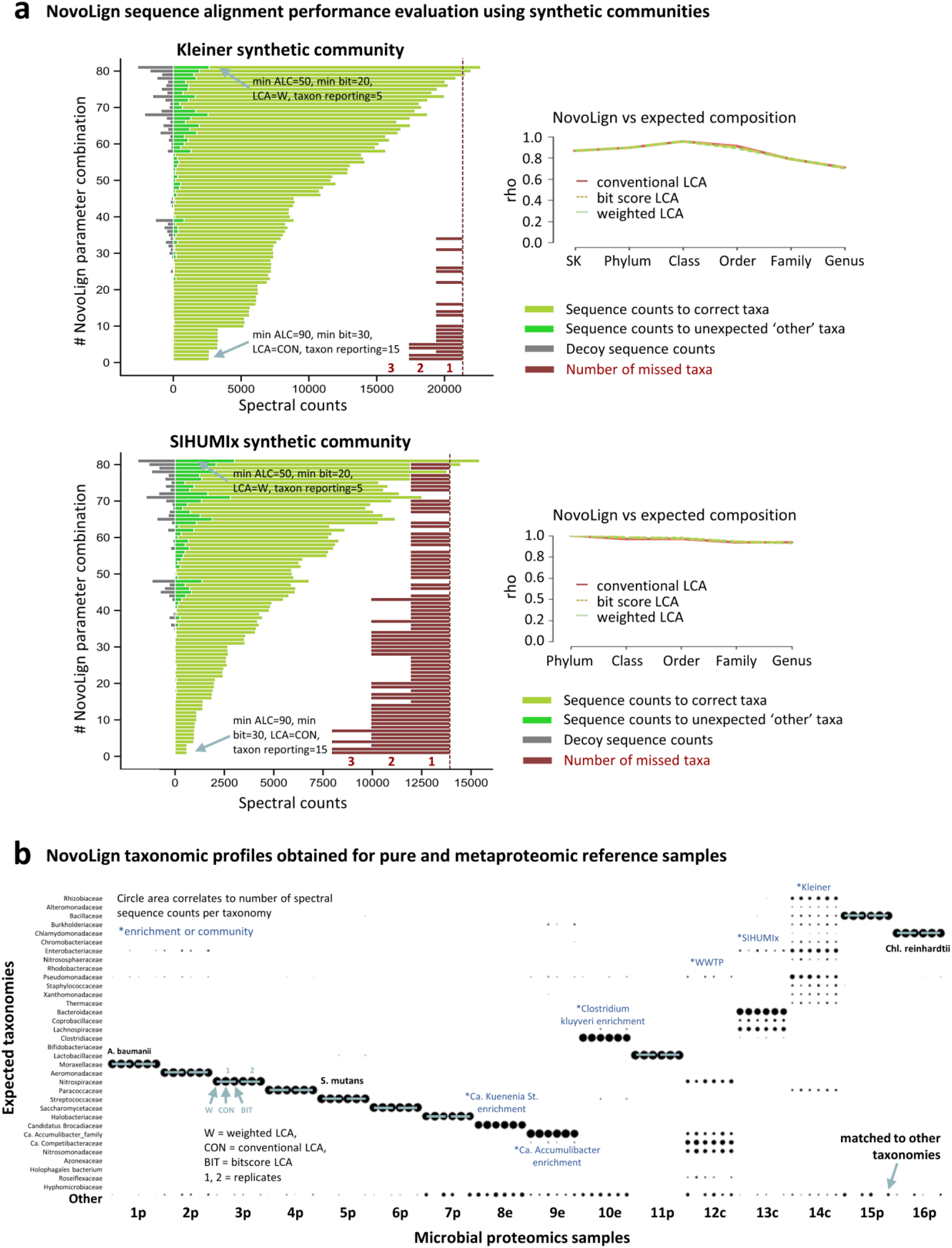
**a)** The left bar graphs depict the NovoLign performance across various parameter combinations (detailed in SI DOC Table 2 and SI EXCEL Table 2) for the synthetic Kleiner equal protein (21 species/strains) and SIHUMIx communities (8 species). The influence of different post-processing parameter combinations, including ALC score, bit-score, LCA, and taxon reporting thresholds, was assessed. The graphs show for each parameter combination the number of i) alignments to expected taxonomies (light green bars), ii) unexpected taxonomies (dark green bars), and the number of iii) decoy sequence matches (grey bars). The red bars located on the right of the bar graphs depict the number of expected taxonomies that were not identified. Generally, the proportion of decoy (random) and other taxonomies were very low for all parameter combinations. Furthermore, missed taxonomies for the Kleiner community were observed only when using very high minimum ALC, bitscore, and taxonomic reporting thresholds. For the SIHUMIx sample, only taxonomies with an abundance <1% were missed. The best parameter combinations (namely those which provided the highest target coverage by maintaining <10% “other” and decoy matches) are listed in SI DOC Tables 3 and 4. The line graphs on the right illustrate the Spearman’s correlation coefficients for both synthetic communities across the taxonomic ranks: superkingdom (SK), phylum, class, order, and family. Both synthetic communities show an excellent correlation between the abundances derived from NovoLign and the expected abundances. For the SIHUMIx sample, the composition determined by database searching served as the reference. Graphs detailing replicate experiments can be found in SI DOC Figure 24. **b)** The figure illustrates the family-level microbial profiles derived from the proteomics and metaproteomics reference samples processed with NovoLign. The size of the circles corresponds to the sequence counts for each taxonomic identifier. Each sample is depicted through six distinct outputs, specifically three different Lowest Common Ancestor (LCA) approaches – weighted LCA (W), conventional LCA (CON), and bitscore LCA (BIT) – for duplicate proteomics samples. The reference samples displayed the expected microbial profiles, with only minor portions of unexpected taxonomies, denoted as “Other” on the y-axis. The labels below the graph (x-axis) provide the microbial sample identifier, with 1p corresponding to *Acinetobacter baumannii*, 2p to *Aeromonas b*., 3p to *Nitrospira moscoviensis*, 4p to *Paracoccus denitrificans*, 5p to *Streptococcus mutans*, 6p to *Saccharomyces cerevisiae*, 7p to *Halanaeroarchaeum sp*., 8e to *Ca*. Kuenenia stuttgartiensis enrichment, 9e to *Ca*. Accumulibacter phosphatis enrichment, 10e to *Clostridium kluyveri* enrichment, 11p to *Lactobacillus saiki*, 12c to aerobic granular sludge community, 13c to SIHUMIx synthetic community, 14c to Kleiner equal protein synthetic community, 15p to *Caldalkalibacillus thermarum*, and 16p to *Chlamydomonas reinhardtii*. The letters next to each number stand for: p = pure reference strain, e = enrichment, and c = community.

**Figure 3.**
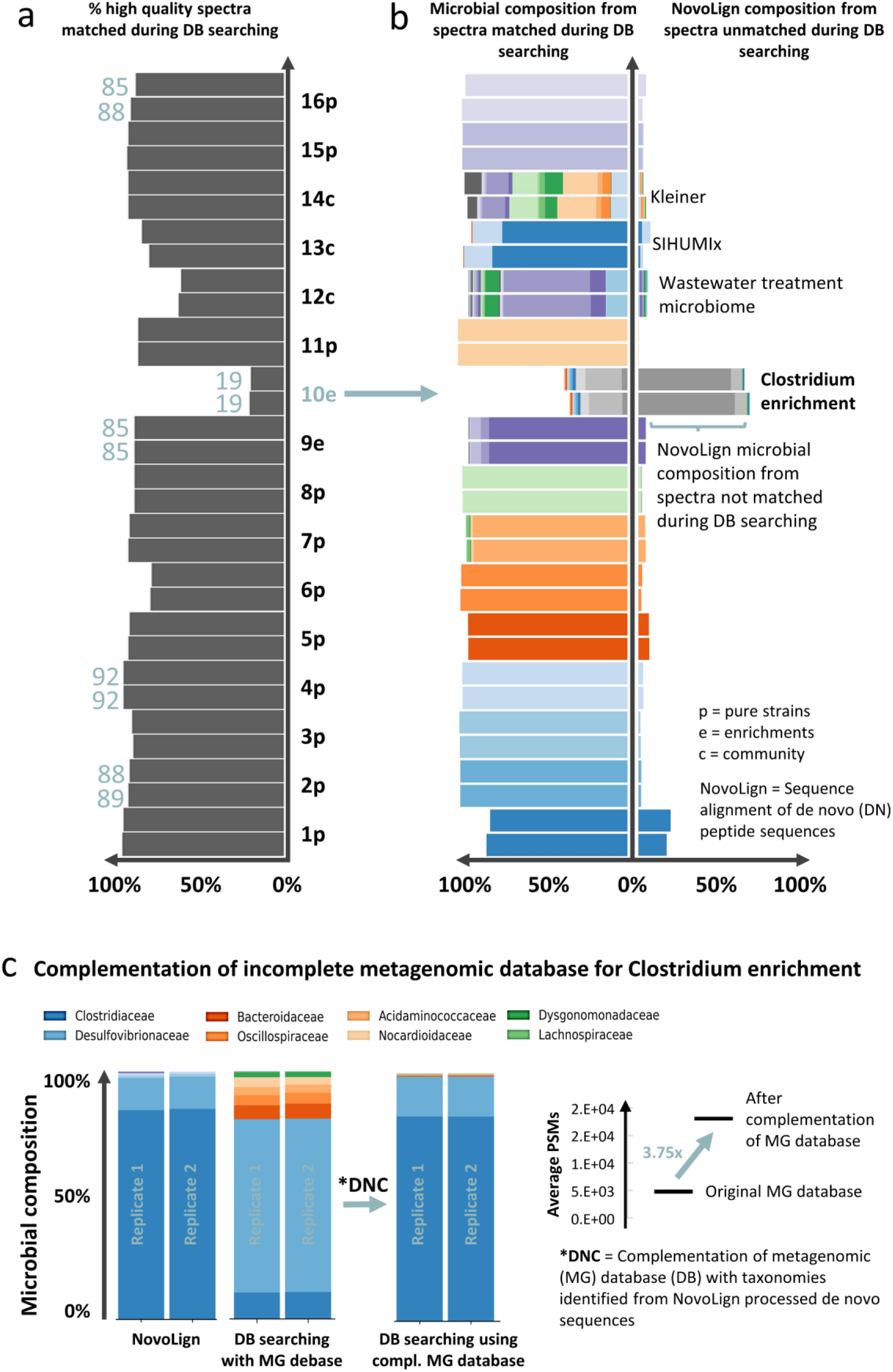
**a)** The graph depicts the percentage of high quality spectra (ALC >90) that were matched during the database searching for the individual (meta)proteomics samples. Except for one sample (*Clostridium kluyveri* enrichment, 10e), the fraction of spectra matched during the database search was very high. The labels next to the bars indicate the microbial sample, with 1p corresponding to *Acinetobacter baumannii*, 2p to *Aeromonas b*., 3p to *Nitrospira moscoviensis*, 4p to *Paracoccus denitrificans*, 5p to *Streptococcus mutans*, 6p to *Saccharomyces cerevisiae*, 7p to *Halanaeroarchaeum sp*., 8e to *Ca*. Kuenenia stuttgartiensis enrichment, 9e to *Ca*. Accumulibacter phosphatis enrichment, 10e to *Clostridium kluyveri* enrichment, 11p to *Lactobacillus saiki*, 12c to aerobic granular sludge microbial community, 13c to SIHUMIx synthetic community, 14c to Kleiner equal protein synthetic community, 15p to *Caldalkalibacillus thermarum*, and 16p to *Chlamydomonas reinhardtii*. The letters next to each number stand for: p = pure reference strain, e = enrichment, and c = community. **b)** The graph compares the taxonomic profiles at the family level obtained by database searching (left bars) with those obtained by processing spectra that were not matched during database searching using NovoLign (right bars). For the majority of the samples, NovoLign provided only a few additional taxonomic annotations, and the taxonomic profiles were very similar compared to those derived from database searching. However, in the case of the Clostridium kluyveri enrichment (10e), NovoLign processing of unmatched spectra showed a substantial number of additional annotations, and a taxonomic profile very different from that obtained through database searching. **c)** The graphs illustrate the complementation of the incomplete metagenomic sequence database for the *Clostridium kluyveri* enrichment (10e). The bars, from left to right, display the microbial composition identified by NovoLign, database searching using the metagenomic database, and database searching using the *de novo* complemented metagenomic database. The taxonomic profile obtained from processing the metaproteomics dataset against the UniRef100 database with NovoLign was significantly different from that obtained through database searching using the metagenomic reference sequence database. Therefore, family level taxonomies identified by NovoLign were extracted from the UniRef100 database and integrated with the metagenomic database (DNC = de novo complementation). The complemented database was then utilized for database searching, providing a profile that closely aligned with the one obtained by NovoLign. Furthermore, the graph on the right highlights the increase in peptide spectrum matches (approx. 3.5 times more compared to using the original metagenomic database) after the database complementation.

### Rapid metaproteomic taxonomic profiling

Rapid taxonomic and functional profiling of complex microbial samples without the need for parallel metagenomics experiments, or extensive databases can be tremendously useful. This allows to rapidly monitor compositional changes over time or when performing quality checks before committing to more extensive experiments, which saves time and experimental costs. As a first step, the sequences are filtered for high quality de novo sequences by applying a minimum *de novo* quality score threshold (e.g. ALC score for PEAKS^64^). Next, the sequences are randomized to provide an additional set of decoy sequences, as already introduced for the NovoBridge pipeline^39^. Both sequence datasets are then aligned, where the decoy alignments provide an estimate for the number of random alignments per dataset. The results are then filtered for confident alignments by applying a minimum bitscore threshold. Furthermore, because sequences often provide alignments to several sequences from different taxonomies, the pipeline includes algorithms to determines a consensus lineage. In order to maximize the annotations at lower taxonomic ranks, we investigated 3 different lowest common ancestor (LCA) approaches. The first “conventional” LCA approach (“CON”) strictly determines the lowest common ancestor from all lineages above the bitscore threshold. The second bitscore approach (“BIT” LCA) is based on the recently published BAT tool^68^ which determines the consensus taxonomy stepwise. Thereby, for every taxonomic rank the taxonomy which accounts for the majority of the total bitscore is chosen. The third approach is a “weighted” approach (“W” LCA), which was introduced for processing of metagenomics data, by Buchfink et al. in 2015^67^. This approach first assigns weights to all taxonomies based on their frequency in the alignment results. Furthermore, the consensus lineage is determined from taxonomies which combined account for at least 80% of the sum of weights to which the query sequence provided alignments. Next, the pipeline performs grouping of the consensus lineages in order to estimate the microbial composition. The sequence counts assigned to each taxonomic group can be seen as an abundance estimate for these organisms. Nevertheless, in order to avoid reporting extensive lists of very low abundant taxonomies (with only few sequence counts) we implemented a minimum frequency threshold for the taxonomic reporting step, as also described earlier^39^.

In order to verify whether the sequence alignment provides an accurate representation of the taxonomies present in the microbial community, we processed two synthetic communities with known content (SI EXCEL Table 1). For this purpose, raw data from the ‘Kleiner equal protein’ and SIHUMIx synthetic communities were de novo sequenced, and processed using NovoLign with various combinations of processing parameters. These included different ALC score, bitscore and taxon reporting thresholds, as well as different LCA approaches. The evaluated parameter combinations are summarized in SI DOC Table S2. The NovoLign processing results are summarized in Figure 2A (and SI DOC Figure 24), where the complete parameter evaluation output is provided in SI EXCEL Table 2. This allowed to evaluate how these parameters impact on the number of i) sequences with correct alignments, ii) sequences with unexpected alignments (to other taxonomies), and the number of iii) aligned decoy sequences. For example, the ALC and bitscore had the largest impact on % other matches and % decoy matches, while the LCA algorithm and the taxonomy reporting threshold mostly influenced the target coverage and the number of identified taxonomies (SI DOC Figures 25–29). The best NovoLign parameter combinations (namely those which provide a high target coverage by maintaining less than 10% “other”, and “decoy matches”) for both synthetic communities, are listed in SI DOC Tables 3 and 4. These generally employed either weighted or bitscore-based LCA, with a minimum ALC of 70 and a minimum bitscore threshold of 25. Nevertheless, although all taxonomies were identified for both synthetic communities at the family level, the families *Bifidobacteriaceae* and *Lactobacillaceae*, present in the SIHUMIx sample, were only observed when employing lower taxonomic reporting thresholds of 5 instead of 15. However, these microbes were also very low abundant in the database searching results, at approx. 0.15 and 0.05%, respectively. Taxonomies with ≥1% relative abundance could all be identified at the family or genus levels. Overall, the microbial composition obtained by these parameters showed a strong correlation to the true (known) composition of both communities, as demonstrated by the strong Spearman’s rank correlation shown in Figure 2A (and SI DOC Figures 30 and 31).

Finally, by using the optimized NovoLign processing parameters, we aimed to demonstrate the application of the NovoLign pipeline using a broad spectrum of taxonomies and sample complexities. Therefore, we *de novo* sequenced and processed a range of pure reference strains, synthetic communities, microbial enrichment cultures, and complex microbial samples using NovoLign (SI EXCEL Table 1). This approach yielded taxonomic profiles for all samples that were very close to the expected (known) microbial composition, or matched those obtained from orthogonal database searching experiments (Figure 2B, and SI DOC Figures 30 and 31).

Compared to the previously developed *de novo* pipeline, which employed exact sequence matches (NovoBridge), sequence alignment significantly increased the number of annotated sequences. For instance, for the *S. cerevisiae, Aeromonas*, and *Nitrospira* reference samples, the bitscore based LCA provided on average a 3.5-fold increase, and the weighted LCA on average an approx. 5-fold increase in annotated sequences at the lower taxonomic levels, order, family and genus (SI DOC Figure 32).

Finally, processing the *S. cerevisiae* dataset (58,368 *de novo* sequences, including decoys) was very time efficient. For example, NovoLign processing using the complete SwissProt database (containing 567,413 sequences, 280 MB) takes only 0.7 minutes, and NovoLign processing with the extensive UniRef100 database (containing 352,965,587 sequences, approx. 180 GB) only 32.4 minutes, respectively (on a desktop with an Intel^(R)^ Core^(TM)^ i7-7700K and 32 GB RAM). A more detailed breakdown of the processing times for the individual NovoLign modules is presented in SI DOC Figure 33.

### Metaproteomic quality plots, database completeness and complementation

A crucial step in metaproteomics is constructing the reference sequence database. To date, there is no consensus on the best method for creating such databases. However, generic reference sequence databases have a very large search space, which reduces sensitivity and increases the likelihood of false discoveries. Additionally, these databases can be regarded as incomplete, potentially missing crucial organisms or proteins^14, 16, 29^. Therefore, constructing sample-specific databases through whole metagenome sequencing is currently regarded as the most effective approach for many metaproteomic applications. However, because metagenomics experiments are time-consuming and expensive, generic databases are often employed to enable rapid quality monitoring in laboratory experiments. Furthermore, several factors can affect the coverage and accuracy of these databases, including DNA extraction, sequencing, genome assembly, identification of open reading frames, and taxonomic classification. Moreover, differences in the timing of sampling and the storage conditions of biomass between metagenomics and metaproteomics experiments can lead to significant discrepancies due to possible degradation and alterations in the microbial composition^14-16, 19, 23^. As a result, constructing an accurate protein sequence database from metagenomic data remains a delicate task.

While the large number of comparable datasets in single-species proteomics allows for setting expectations for peptide spectrum matches, the increased complexity of metaproteomics samples makes it more difficult to monitor the quality of these experiments. Consequently, poorly constructed reference sequence databases that miss members of the community may go unnoticed without the application of additional approaches.

One advantage of the alternative *de novo* sequencing in proteomics is that it provides next to a likely amino acid sequence also a quality metric for each spectrum. Spectra with good fragment ion coverage will obtain a high quality score, and are also expected to give a strong peptide spectrum match if the sequence is present in the reference database.

This offers the opportunity for NovoLign to identify missed taxonomies or proteins through processing *de novo* sequences from unmatched high-quality spectra. In order to demonstrate this procedure we determined the fraction of high-quality spectra (ALC≥90) that were matched during database searching for our employed reference strains, enrichment cultures, synthetic and natural communities (Figure 3A). In order to investigate for missed taxonomies we also processed the *de novo* sequences from unmatched high-quality spectra through NovoLign and compared the taxonomic profile with those obtained from database searching (Figure 3B).

Interestingly, for most samples more than 80% of the high-quality spectra were matched during database searching, with a strong agreement between the taxonomic profiles obtained by NovoLign and database searching. The remaining fraction of unmatched high-quality spectra did also not reveal new organisms for most samples, and may therefore originate from modified peptides that were not considered during the database search process.

However, in one sample (“*Clostridium kluyveri* enrichment”), the fraction of matched high-quality spectra was less than 25% (Figure 3A, 10e), and the taxonomic profile obtained from NovoLign significantly differed from the one obtained through database searching (Figures 3B and 3C). For instance, while the NovoLign composition indicates that the family *Clostridiaceae* is dominant (approximately 80%), the outcomes from database searching shows that *Desulfovibrionaceae* is dominant, with *Clostridiaceae* being only a minor component. This reveals that the metagenomic reference database used for database searching poorly represents the sample analyzed in the proteomics experiment. Although there could be several reasons for this (as described above), the observed discrepancy is most likely due to differences in storage times and sample processing between metagenomics and metaproteomics experiments.

Advantageously, NovoLign allows also to extract sequences of the identified taxonomies from the UniRef100 database. These can then be used to complement the existing database with the missing organisms. Therefore, we complemented the original *Clostridium kluyveri* enrichment database with the *de novo* identified taxonomies and repeated the database searching with the complemented database. This significantly increased the number of peptide-spectrum matches (3.75x compared to the original metagenomic database) and aligned the taxonomic profiles with the one obtained by NovoLign (Figure 3C). The microbial profile obtained after database complementation is also consistent with the 16S rRNA amplicon sequencing data and performed reactor experiments for the same enrichment previously^60^.

## CONCLUSIONS

Here, we introduce a novel *de novo* metaproteomics pipeline based on sequence alignment, called NovoLign. This pipeline enables rapid taxonomic profiling with deep coverage and allows for the evaluation of the quality of metaproteomics experiments and reference sequence databases. Additionally, NovoLign facilitates the complementation of reference databases with sequences from organisms not present in the database.

We optimized the DIAMOND alignment using a wide array of in silico peptide data, which represent common errors found in *de novo* sequencing. Moreover, we assessed the NovoLign pipeline with a broad spectrum of pure reference strains, synthetic communities, laboratory enrichment cultures, and environmental microbial communities. We evaluated post-processing parameters, including taxonomic grouping, to enhance taxonomic coverage and minimize false positive annotations. Finally, the taxonomic profiles obtained for the reference strains and synthetic communities closely matched the expected taxonomic composition up to the family and genus levels, with less than 10% unexpected taxonomies or decoy sequence matches. Complete proteomics datasets (# 50K *de novo* sequences) can be aligned to Swiss-Prot in less than one minute and UniRef100 to approx. 30 minutes, using a conventional desktop PC. While earlier tools only used exact matches and precomputed peptide databases, the flexibility of NovoLign allows to employ custom reference sequence databases and variable proteolytic cleavage sites, as well as annotation of organisms not present in the database to their closest taxonomic neighbor.

Furthermore, the combination of *de novo* sequencing with sequence alignment allows to identify fragmentation spectra of modified peptides, which otherwise remain unmatched in complex metaproteomics experiments^61, 70, 71^. Finally, advanced *de novo* (meta)proteomics pipelines will become increasingly important in many fields of research where obtaining reference sequence databases is challenging or not possible at all. For example, *de novo* sequence alignment pipelines can be useful for assembling antibody sequences in cases where cloning and sequencing of the coding mRNA is not possible^36, 37^, or for the identification of HLA-associated peptides that derive from modified or mutated sequences, spliced peptides generated in the proteasomes, or from non-coding regions^72^. Additionally, viruses are increasingly studied in the context of bacteriophages which can selectively target and kill bacteria. However, many viruses mutate with high frequency, which makes their proteomic analysis complicated using conventional database searching approaches^73^.

In summary, NovoLign enables rapid taxonomic profiling using sequence alignment and serves as an orthogonal approach for assessing the quality of metaproteomics experiments that use database searching. The pipeline furthermore supports complementation of reference sequence databases to improve sequence coverage during database searching.

## Supporting information

Kleikamp_et_al_2024_NovoLign_SI_DOC

Kleikamp_et_al_2024_SI_EXCEL_Table_1

Kleikamp_et_al_2024_SI_EXCEL_Table_2

## Data availability

The mass spectrometry proteomics raw data acquired within this project have been deposited in the ProteomeXchange consortium database with the dataset identifier PXD050548. ProteomeXchange identified for all other datasets are provided in the methods section. Correspondence and requests for materials should be addressed to HK and MPa.

## Code availability

The NovoLign pipeline is publicly available via: https://github.com/hbckleikamp/NovoLign.

### User contribution

Author contributions. HK and MPa conceived and designed the project. HK, MPa, RZ, RV, MPr, PS, JvE and MA performed the experiments. HK, MPa, RZ, RV, MvL provided critical input to experiments and/or protocols. HK and MP conducted the data analysis. HK and MPa generated the figures and wrote the original draft. All coauthors contributed in reviewing and editing the manuscript. ^#^Note: RvZ and RvV contributed equally to this work.

## Acknowledgements

The authors acknowledge Dita Heikens for her support with sample preparation and all other colleagues from the Department of Biotechnology for valuable discussions. This work was supported by a TU Delft Startup fund and the SIAM Gravitation Grant 024.002.00, the Netherlands Organization for Scientific Research (NWO).

## Competing interests

The authors declare that they have no known competing financial interests or personal relationships that could have appeared to influence the work reported in this paper.

## Notes

### Competing Interest Statement

The authors have declared no competing interest.

