## Supplementary material for "NovoLign: metaproteomics by sequence alignment": Kleikamp_et_al_2024_NovoLign_SI_DOC

### Table of Contents

| Section | Page |
|---|---|
| A. Construction of a synthetic in silico de novo sequence dataset | 2 |
| B. Optimizing Diamond sequence alignment parameters | 4 |
| *B1. Scoring matrices, seed shapes and seed search algorithm* | |
| *B2. Query cover and percentage of sequence identity* | |
| *B3. Gap penalties* | |
| C. Sequence alignment of different de novo error types and mutations | 8 |
| D. Evaluation of NovoLign pipeline for performance, coverage and % false positives | 18 |
| *D1. Overview of obtained sequence counts and abundance correlation for the parameter combinations.* | |
| *D2. Impact of ALC, bit and cutoff thresholds on % decoy and % other matches* | |
| *D3. Impact of LCA algorithm on % decoy, % other matches and total taxon counts* | |
| *D4. Influence of various LCA algorithms and bit scores on total sequence* | |
| *D5. Overview of best-performing NovoLign parameters for the synthetic equal protein "Kleiner community"* | |
| *D6. Overview of best-performing NovoLign parameters for the synthetic "SIHUMIx community"* | |
| *D7. Abundance profiles for the synthetic "equal protein community" for different LCA algorithms and taxonomic frequency cutoffs* | |
| *D8. Abundance profiles for the synthetic "SIHUMIx community" for different LCA algorithms and taxonomic frequency cutoffs* |  |
| *D9. Comparison of sequence coverage between NovoLign and NovoBridge* | |
| *D10. Processing time performance of NovoLign pipeline* | |
| E. SI References | 30 |

### A. Construction of a synthetic in silico de novo sequence dataset

We developed a code ("de novo error-SIM") which constructs reference sequence datasets that not only resemble the common peptide lengths, but also common de novo sequencing errors and mutated peptides (SI Figure 1). The python codes and reference sequence data sets used in this study are publicly available via: https://github.com/hbckleikamp/De-Novo-ErrorSIM. The script “mutate_swap.py” simulates de novo errors by mutation, substitution, or inversion, based on the error classes described by Muth & Renard (2018)[1]. The “Levenshtein.py” script calculates the Levenshtein distance for each error sequence to select the desired error rates within datasets. The “error_combination.py” script combines different types of errors based on observed frequency in Muth et al., 2018[1], into single reference datasets. To establish a realistic de novo error dataset, the chance of a specific error occurring per peptide length was determined. The longer the peptide, the more likely this peptide will contain more than 1 error, and the more likely the peptide will be misannotated. Therefore, the Levenshtein distance to the original peptide was calculated[2]. Each error type (i.e. mutations, inversions and substitutions) shows a different Levenshtein distance. A mutation has a smaller effect on the Levenshtein distance, and could in theory occur in each amino acid. Inversions can occur only once every amino acid segment (e.g. consisting of 2–3 amino acids), but these errors have a larger effect on the distance. Substitutions of equal mass (e.g. 2 to 3 amino acids) can only occur for specific combinations of amino acids within one peptide. Therefore, to achieve similarly “different” peptides across different de novo error datasets, the rate of occurrence within peptides are important factors to consider. Each error type was simulated at different rate of occurrence generate sets with similar Levenshtein distance to a 5% mutation chance.

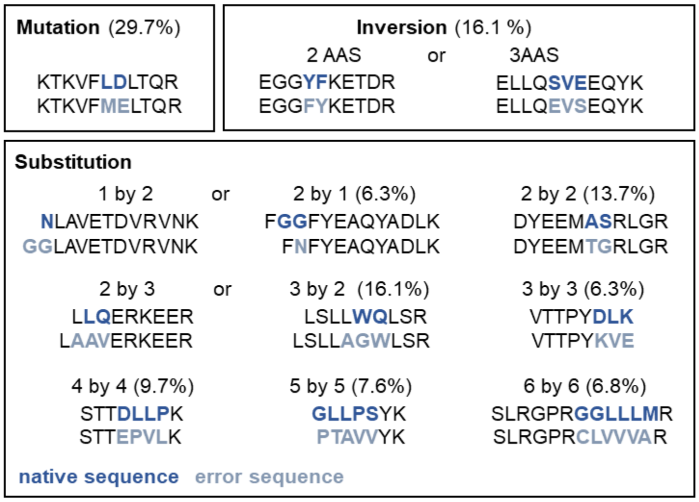


**Mixed DN error dataset**

| Error class | rate (%) |
|---|---|
| Substitution of 1 by 1 or 2 AAs | 6.3 |
| Substitution of 2 by 2 AAs | 13.7 |
| Substitution of 3 by 3 AAs | 6.3 |
| Inversion of 2 or 3 AAs | 16.1 |
| Substitution of 2 by 3 AAs | 3.7 |
| Substitution of 4 by 4 AAs | 9.7 |
| Substitution of 5 by 5 AAs | 7.6 |
| Substitution of 6 by 6 AAs | 6.8 |
| Mutation | 29.7 |

**SI Figure 1.** The graph shows examples for the 12 de novo error classes based on mutation, inversion and substitution. The table on the left details the combined dataset. The fraction of each error is based on experimental data. Each dataset contains 1000 sequences (8–50 AA length).

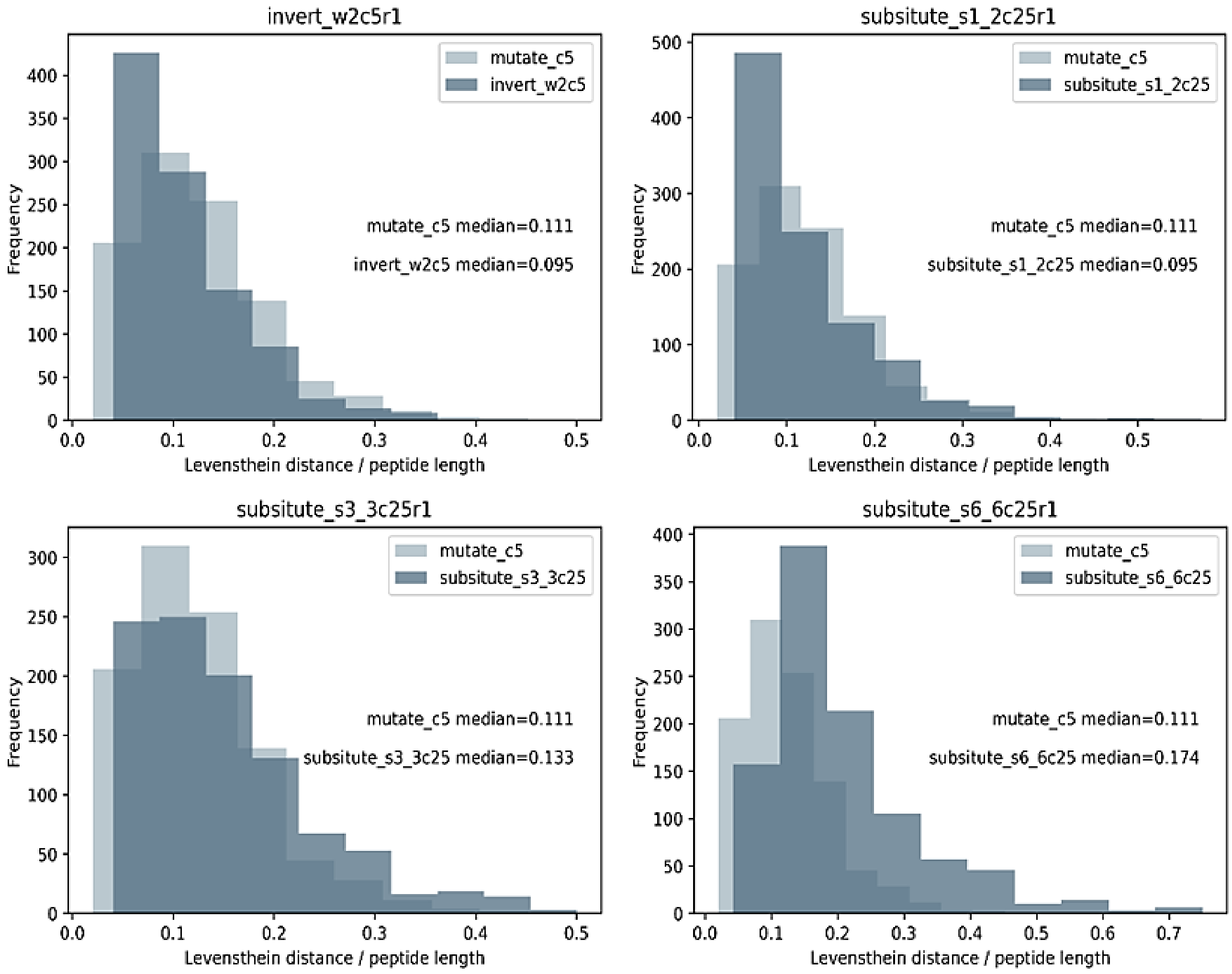


**SI Figure 2.** Frequency versus Levenshtein distance over peptide length for different error types at a mutation rate of 5%. The Levenshtein distance is the minimum number of operations needed to transform sequence A into sequence B , where the only allowed operations are substitution, deletion or insertion of a single character.

### B. Optimizing Diamond sequence alignment parameters

We evaluated the performance of scoring matrices, seed shapes, search algorithms for seeds, gap penalties, and the reporting parameters query coverage and percentage of sequence identity. The performance was expressed as F1 scores for precision and recall, and the length distribution of aligned sequences was visualized by violin plots. For this initial parameter evaluation, the minimum bit score was set to 20, for reporting of target sequences -k was set to 50, and the effective database (--dbsize) was set to 1 (to overcome too rigid e-value cutoffs). Additionally, we employed the default (fast) mode, which is more efficient for short reads. The sensitive modes are recommended for longer sequences (--sensitive or --more-sensitive).[3]

#### B1. Scoring matrices, seed shapes and seed search algorithm

The default substitution matrix used by DIAMOND is BLOSUM62 ("BLOcks SUbstitution Matrix", 62 = built using sequences with less than 62% similarity)[3]. This matrix is commonly used for sequence alignment between evolutionally divergent protein sequences. It considers highly conserved regions in series of alignments forbidden to contain gaps[4]. Other widely employed scoring matrices are the PAM "Point Accepted Mutation" matrices (higher numbers in the PAM matrix denote larger evolutionary distance), which are based on mutations observed throughout a global alignment, including both highly conserved as well as highly mutable regions. While the PAM matrices score all amino acid positions in related sequences, the BLOSUM matrices are based on substitutions and conserved positions in blocks. This makes the PAM matrices in general more suitable for the alignment of short peptide sequences.[4]

In our study, we evaluated the scoring and reporting parameters for DIAMOND, but we did not investigate a modified scoring matrix. Nevertheless, because leucine and isoleucine cannot be distinguished in conventional proteomics experiments, we modified all databases used with DIAMOND by replacing "I" by "L".

DIAMOND also employs longer spaced seeds to accelerate the searching process. The default sensitivity mode uses seeds with a length of 12 and 15, respectively, which limits the alignment of shorter sequences. Furthermore, DIAMOND offers 3 seed search algorithms --algo (0,1,ctg), where 0 means double-indexed (designed for large input files but less efficient for small query files), 1 means query-indexed (improves performance for small query files), and contiguous-seed mode (ctg) which further improves performance for shorter sequences and small query files[3]. We evaluated therefore a matrix of conditions using the combined de novo sequence error dataset "combined_r1.fasta" and the Swiss-Prot (569,793 entries, 23 July 2023) and UniRef100 (356,800,925 entries, 23 July 2023) databases, combining different seeds, algorithms and scoring matrices (Figures 3–4). Considering the f1 score and peptide length distribution (in particular alignment of short sequences), the PAM30 and PAM70 matrices generally performed better compared to BLOSUM62. Albeit the shortest seeds with the seed search algorithm setting 0 and 1 performed best, the default seed in contiguous-seed mode performed comparably well.

```
seeds=[('1111','0'),
       ('1111','1'),
       ('1111','ctg'),
       ('111111','0'),
       ('111111','1'),
       ('111111','ctg'),
       ('Default','0'),
       ('Default','1'),
       ('Default','ctg')]

matrices=["PAM30","PAM70","BLOSUM62"]
```

```
command="".join(['"'+diamond_path+'"',
        " blastp -q " +'"'+query_path+'"',
        " -d "+'"'+db_path+'"',
        " -o "+'"'+output_+'"',
        " -c1 -b 1 ",
        " -k50 ",
        " --log ",
        " --matrix "+matrix,
        " --shape-mask "+seed,
        " --algo "+algr,
        " --dbsize 1 ",
        " -f 6 qseqid sseqid stitle pident bitscore evalue qseq sseq full_sseq ",
        " -t "+'"'+diamond_folder+'"'])
```

**SI Figure 3.** DIAMOND commands for comparing combinations of different matrices ("matrix"), seeds ("seed") and seed search algorithms (--algo).

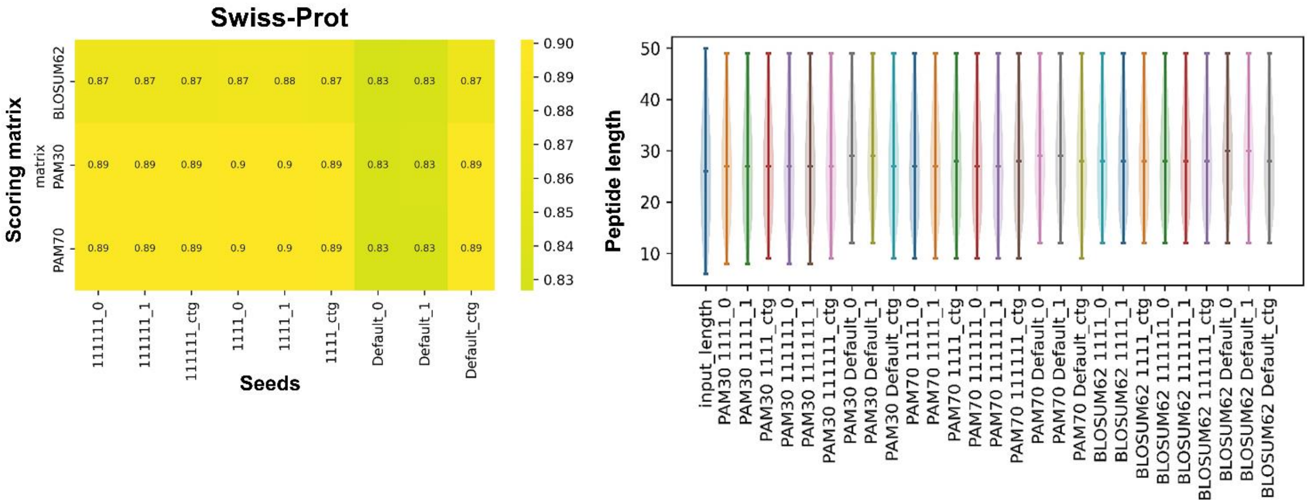


**SI Figure 4.** Evaluation of scoring matrices and seeds (seed shape and algorithm pairs, right F1 score matrix) as well as length distribution of aligned peptides (left, violin plots). The default DIAMOND settings operate with seeds that scan for longer sequences, and shorter peptides are therefore not aligned. For example, when using default parameters (e.g. PAM30 or PAM70, default seeds and seed search algorithm "0" or "1") no peptides below the length 12 were aligned. However, shorter peptides (e.g. 8 amino acids) were aligned when using the shorter custom seeds as well as when using default seeds in combination with the contiguous seed search algorithm (right, violin plot). These conditions provide also the best F1 score (left, scoring matrix). Generally, the PAM matrices showed a better F1 score as well as amino acid length distribution compared to BLOSOM62.

### B2. Gap penalties

Standard scoring matrixes commonly use a high gap opening penalty and low gap extension penalties. For example, the default parameters in DIAMOND for --gapopen (gap open penalty) and --gapextend (gap extension penalty) are 11 and 1, respectively. This heavily penalizes gaps in general, but enables more distant alignments with larger gaps, which is disfavoring the alignment of de novo error sequences. As described earlier, common errors include unequal substitutions of amino acids, such as 1 to 2 or 2 to 3 amino acids, which result in small gaps. Similarly, inversion of 2 or 3 amino acids could result in a small, gapped alignment. Therefore, the gap penalty should rather be low. On the other hand, to prevent spurious alignments, the gap extension penalty may rather be increased. In order test our hypothesis, we evaluated different gap opening and gap extension penalty combinations.

```
for c0 in [0,2,4,6,8,10]:       # gap opening
    for c1 in [0,2,4,6,8,10]:   # gap extentions

        query_path=files[r]
        qfilename=files[r]

        output_="output_placeholder"


        command="cd "+'"'+basedir +'"'+ " & " + \
                "".join(['"'+diamond_path+'"',
                " blastp -q " +'"'+query_path+'"',
                " -d "+'"'+db_path+'"',
                " -o "+'"'+output_+'"',
                " -c1 -b 1 ",
                " -k50 ",
                " --log ",
                " --id 85 ",
                " --query-cover 85 ",
                " --custom-matrix "+'"'+matrix_path+'"'+" --gapopen "+str(c0)+"  --gapextend "+str(c1)+" ",
                " --dbsize 1 --algo ctg"
                " -f 6 qseqid sseqid stitle pident bitscore evalue qseq sseq full_sseq ",
                " -t "+'"'+diamond_folder+'"'])
```

**SI Figure 5.** DIAMOND commands for comparing different gap opening (--gapopen) and gap extension (--gapextend) combinations. The survey used in addition the earlier identified parameters for query cover and % identity, as well default seeds (sensitivity mode) plus the contiguous seed search algorithm.

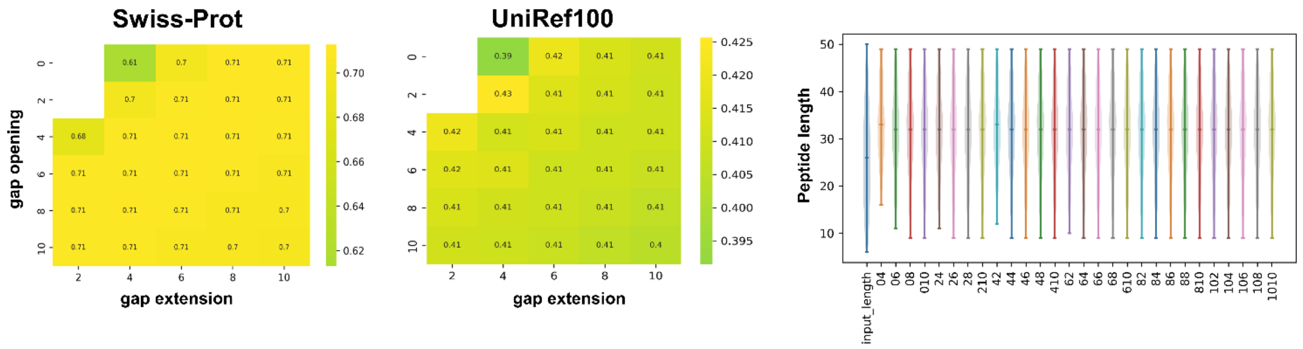


**SI Figure 6.** Evaluation of the parameters gap opening and extension for the PAM70 matrix aligned to the Swiss-Prot or the UniRef100 database (right, F1 scoring matrix) as well as length distribution of the aligned peptides (left, violin plots). Along the x-axis of the violin plot, combinations of gap opening and gap extension penalties are plotted. The first number indicates the gap opening penalty, while the second denotes the gap extension penalty. For example, '06' represents a gap opening penalty of 0 and a gap extension penalty of 6. The first plot “input_length” shows the original peptide length distribution of the input dataset. The Swiss-Prot alignment showed a range of combinations that performed equally well, however, the UniRef100 alignment showed that a combination of gap opening / gap extension = 2/4 provides the best F1 score (including a broad peptide length distribution).

#### B3. Query cover and percentage of sequence identity

The reporting parameters query cover and percentage of sequence identity can have a significant impact on the number of recalls and precision. Therefore, we evaluated for the best combination between query coverage and percentage of sequence identity for short de novo peptides.

```
for c1 in [75,80,85,90]:
```

```
output_="output_placeholder"
command="cd "+'"'+basedir +'"'+ " & " + \
        "".join(['"'+diamond_path+'"',
        " blastp -q " +'"'+query_path+'"',
        " -d "+'"'+db_path+'"',
        " -o "+'"'+output_+'"',
        " -c1 -b 1 ",
        " -k50 ",
        " --log ",
        " --id 75 ",
        " --query-cover  "+str(c1)+" ",
        " --matrix "+matrix,
        " --dbsize 1 ",
        " -f 6 qseqid sseqid stitle pident bitscore evalue qseq sseq full_sseq ",
        " -t "+'"'+diamond_folder+'"'])
```

```
for c0 in [75,80,85,90]:
```

**SI Figure 7.** DIAMOND commands for comparing different percent identity (--id) and coverage (–query-cover) combinations.

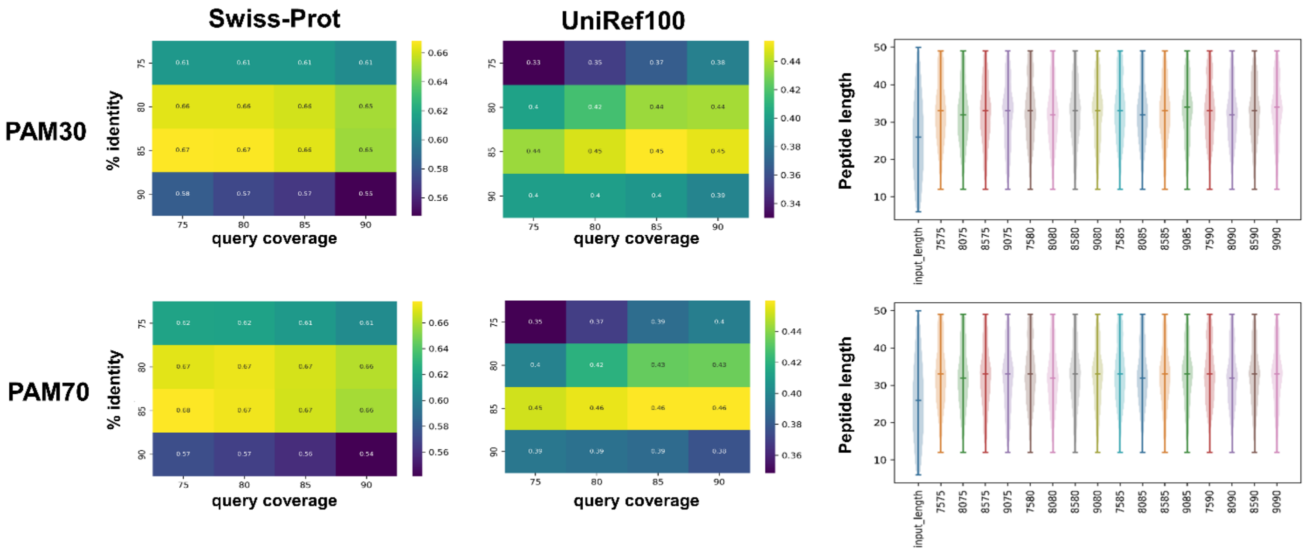


**SI Figure 8.** Evaluation of the reporting parameters % identity and query coverage (right, F1 matrix) as well as length distribution of aligned peptides (left, violin plots), for PAM30 and PAM70, aligned to Swiss-Prot as well as UniRef100. Along the x-axis of the violin plot, combinations of percent identity and coverage are plotted. The first two numbers indicate the percent identity, while the second two denote the percent coverage. For example, '7575' represents a percent identity of 75 and a percent coverage of 75. The first plot "input_length" shows the original peptide length distribution of the input dataset. PAM70 performs slightly better compared to PAM30, with the best combination % identity (--id) 85 and query coverage (--query-cover) 75.

### C. Sequence alignment of different de novo error types and mutations

In order to provide a general overview of the ability to correctly align sequences with different de novo sequencing errors and mutations, we performed a series of alignments for the different de novo error types individually (using parameter combinations identified in the previous experiments). Table 1 summarizes the different de novo error datasets. Thereby, all sequences were aligned to the UniRef100 database and analyzed for % alignments which provide an exact (sequence) match, correct (taxonomy) match, false-positive match, random matches, and unmatched fraction. Finally, the error length distribution of the aligned peptides was compared to the length distribution of the input dataset.

**SI Table 1.** Table of generated and analyzed de novo error reference datasets. Each dataset contains 1000 modified peptide sequences retrieved from Swiss-Prot entries.

| # | File | Error simulation |
|---|---|---|
| 1 | combined_r1.fasta | replicate 1 of combined de novo errors (r1 files) according to observed error rates (Muth et al., 2018) |
| 2 | combined_r2.fasta | replicate 1 of combined de novo errors (r1 files) according to observed error rates (Muth et al., 2018) |
| 3 | combined_r3.fasta | replicate 1 of combined de novo errors (r1 files) according to observed error rates (Muth et al., 2018) |
| 4 | invert_w2c10r1.fasta | Random shuffle of 2 consecutive amino acids at a 10% occurrance chance |
| 5 | invert_w2c1r1.fasta | Random shuffle of 2 consecutive amino acids at a 1% occurrance chance |
| 6 | invert_w2c20r1.fasta | Random shuffle of 2 consecutive amino acids at a 20% chance |
| 7 | invert_w2c5r1.fasta | Random shuffle of 2 consecutive amino acids at a 5% chance |
| 8 | invert_w3c10r1.fasta | Random shuffle of 3 consecutive amino acids at a 10% chance |
| 9 | invert_w3c1r1.fasta | Random shuffle of 3 consecutive amino acids at a 1% chance |
| 10 | invert_w3c20r1.fasta | Random shuffle of 3 consecutive amino acids at a 20% chance |
| 11 | invert_w3c5r1.fasta | Random shuffle of 3 consecutive amino acids at a 5%chance |
| 12 | mutate_c10r1.fasta | Randomly mutate reverse translated amino acids at 10% chance |
| 13 | mutate_c1r1.fasta | Randomly mutate reverse translated amino acids at 1% chance |
| 14 | mutate_c20r1.fasta | Randomly mutate reverse translated amino acids at 20% chance |
| 15 | mutate_c5r1.fasta | Randomly mutate reverse translated amino acids at 5% chance |
| 16 | subsitute_s1_2c100r1.fasta | Randomly substitute equal 1 to 2 amino acid mass pairs at 100% chance |
| 17 | subsitute_s1_2c25r1.fasta | Randomly substitute equal 1 to 2 amino acid mass pairs at 25% chance |
| 18 | subsitute_s1_2c50r1.fasta | Randomly substitute equal 1 to 2 amino acid mass pairs at 50% chance |
| 19 | subsitute_s2_1c100r1.fasta | Randomly substitute equal 2 to 1 amino acid mass pairs at 100% chance |
| 20 | subsitute_s2_1c25r1.fasta | Randomly substitute equal 2 to 1 amino acid mass pairs at 25% chance |
| 21 | subsitute_s2_1c50r1.fasta | Randomly substitute equal 2 to 1 amino acid mass pairs at 50% chance |
| 22 | subsitute_s2_2c100r1.fasta | Randomly substitute equal 2 to 2 amino acid mass pairs at 100% chance |
| 23 | subsitute_s2_2c25r1.fasta | Randomly substitute equal 2 to 2 amino acid mass pairs at 25% chance |
| 24 | subsitute_s2_2c50r1.fasta | Randomly substitute equal 2 to 2 amino acid mass pairs at 50% chance |
| 25 | subsitute_s2_3c100r1.fasta | Randomly substitute equal 2 to 3 amino acid mass pairs at 100% chance |
| 26 | subsitute_s2_3c25r1.fasta | Randomly substitute equal 2 to 3 amino acid mass pairs at 25% chance |
| 27 | subsitute_s2_3c50r1.fasta | Randomly substitute equal 2 to 3 amino acid mass pairs at 50% chance |
| 28 | subsitute_s3_2c100r1.fasta | Randomly substitute equal 3 to 2 amino acid mass pairs at 100% chance |
| 29 | subsitute_s3_2c25r1.fasta | Randomly substitute equal 3 to 2 amino acid mass pairs at 25% chance |
| 30 | subsitute_s3_2c50r1.fasta | Randomly substitute equal 3 to 2 amino acid mass pairs at 50% chance |
| 31 | subsitute_s3_3c100r1.fasta | Randomly substitute equal 3 to 3 amino acid mass pairs at 100% chance |
| 32 | subsitute_s3_3c25r1.fasta | Randomly substitute equal 3 to 3 amino acid mass pairs at 25% chance |
| 33 | subsitute_s3_3c50r1.fasta | Randomly substitute equal 3 to 3 amino acid mass pairs at 50% chance |
| 34 | subsitute_s4_4c100r1.fasta | Randomly substitute equal 4 to 4 amino acid mass pairs at 100% chance |
| 35 | subsitute_s4_4c25r1.fasta | Randomly substitute equal 4 to 4 amino acid mass pairs at 25% chance |
| 36 | subsitute_s4_4c50r1.fasta | Randomly substitute equal 4 to 4 amino acid mass pairs at 50% chance |
| 37 | subsitute_s5_5c100r1.fasta | Randomly substitute equal 5 to 5 amino acid mass pairs at 100% chance |
| 38 | subsitute_s5_5c25r1.fasta | Randomly substitute equal 5 to 5 amino acid mass pairs at 25% chance |
| 39 | subsitute_s5_5c50r1.fasta | Randomly substitute equal 5 to 5 amino acid mass pairs at 50% chance |
| 40 | subsitute_s6_6c100r1.fasta | Randomly substitute equal 6 to 6 amino acid mass pairs at 100% chance |
| 41 | subsitute_s6_6c25r1.fasta | Randomly substitute equal 6 to 6 amino acid mass pairs at 25% chance |
| 42 | subsitute_s6_6c50r1.fasta | Randomly substitute equal 6 to 6 amino acid mass pairs at 50% chance |

When evaluating the obtained alignments, the number of correct matches (=same taxonomy) and exact matches (match to original peptide sequences, and taxonomy), were very high for Swiss-Prot and UniRef100 alignments. For example, at the genus level, Swiss-Prot provided >60% successful alignments and the significantly larger UniRef100 database approx. 50%. Most importantly, the number of false matches was very low for all databases, experiments, and taxonomic ranks (<10%), except for the phylum level. Furthermore, the length distribution of peptide sequences matched well the input length distribution. Interestingly, albeit using the default DIAMOND seed (sensitivity mode) even peptides shorter than 10 AA provided correct alignments.

UniRef100 alignments, however, obtained a larger fraction of random (=decoy database) matches, which can be filtered out by increasing the bit score cutoff. However, randomly mutated, reverse translated amino acids at (e.g. at 20% chance) did only provide very poor alignments. Similarly challenging were 3 to 3 amino acid substitutions at 100% chance.

```
def align(input_files,Output_directory,
                      matrix,
                      seed,
                      database_path,
                      diamond_path=str(Path(basedir,"diamond")),
                      output_columns=["qseqid","sseqid","stitle","pident","bitscore","qseq","sseq"],
                      select=" -k50 ",
                      block_size=5,
                      index_chunks=1,
                      minimum_pident=80,
                      minimum_coverage=80,
                      minimum_bitscore=20,
                      gap_open=0,
                      gap_extend=8
                      ):
```

```
command="cd "+'"'+basedir +'"'+ " && " + \
        "".join(['"'+diamond_path+'"',
        " blastp -q " +'"'+input_file+'"',
        " -d "+'"'+database_path+'"',
        " -o "+'"'+output_file+'"',
        " -c" + str(index_chunks),
        " -b "+ str(block_size),
        " "+select+" ",
        " --log ",
        " --custom-matrix "+'"'+matrix_path+'"'+" --gapopen "+str(gap_open)+" --gapextend "+str(gap_extend),
        " --algo ctg --dbsize 1 ",
        " --id "+          str(minimum_pident),
        " --min-score "+   str(minimum_bitscore),
        " --query-cover  "+str(minimum_coverage),
        " -f 6 qseqid "+" ".join(output_columns)+" ",
        " -t "+'"'+Temporary_directory+'"'])
```

**SI Figure 9.** DIAMOND command for aligning individual de novo error datasets (as outlined in SI Table 1) to Swiss-Prot and UniRef100. The selection of individual parameters is based on the evaluation experiments described above. The employed scoring matrix was throughout the experiments shown in the following PAM70.

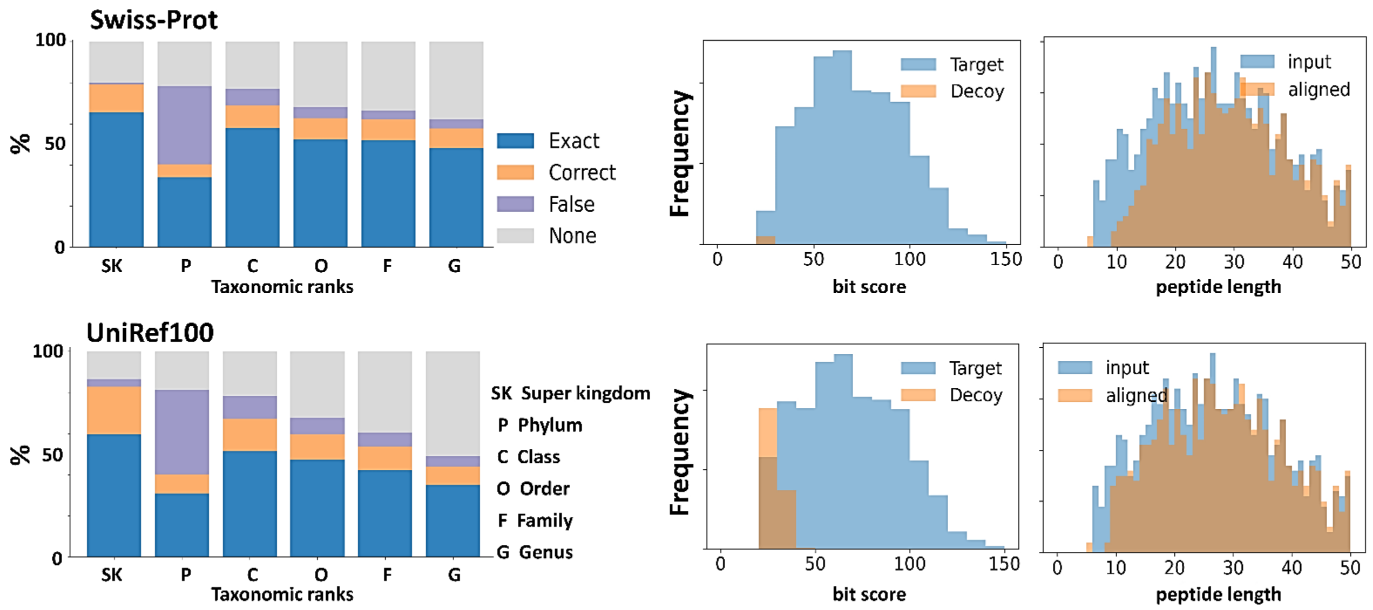


**SI Figure 10.** Sequence alignment recall, precision, random and false matches, bit score and peptide length distribution for combined de novo error dataset #1 (combined_r1.fasta) which covers all types of de novo errors (SI Table 1) with a composition frequency according to previously observed error rates by Muth et al., 2018. Sequences were aligned to the Swiss-Prot and UnRef100 databases.

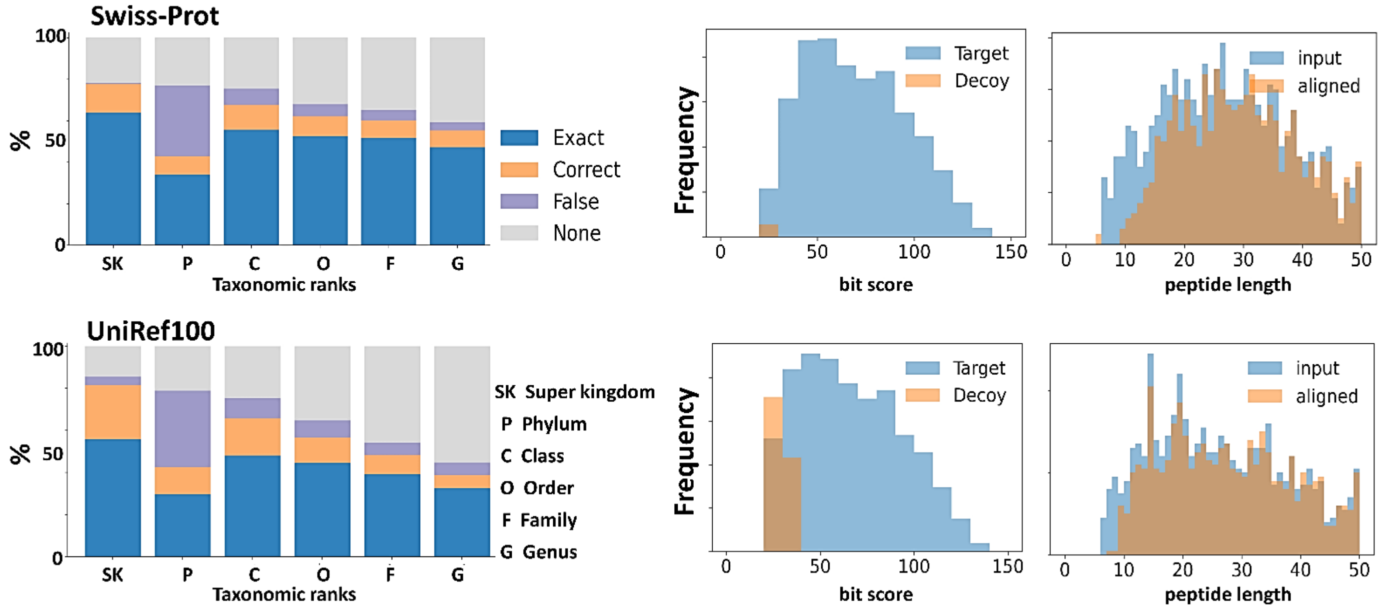


**SI Figure 11.** Sequence alignment recall, precision, random and false matches, bit score and peptide length distribution for combined de novo error dataset #2 (combined_r2.fasta) which covers all types of de novo errors (SI Table 1) with a composition frequency according to previously observed error rates by Muth et al., 2018. Sequences were aligned to the Swiss-Prot and UnRef100 databases.

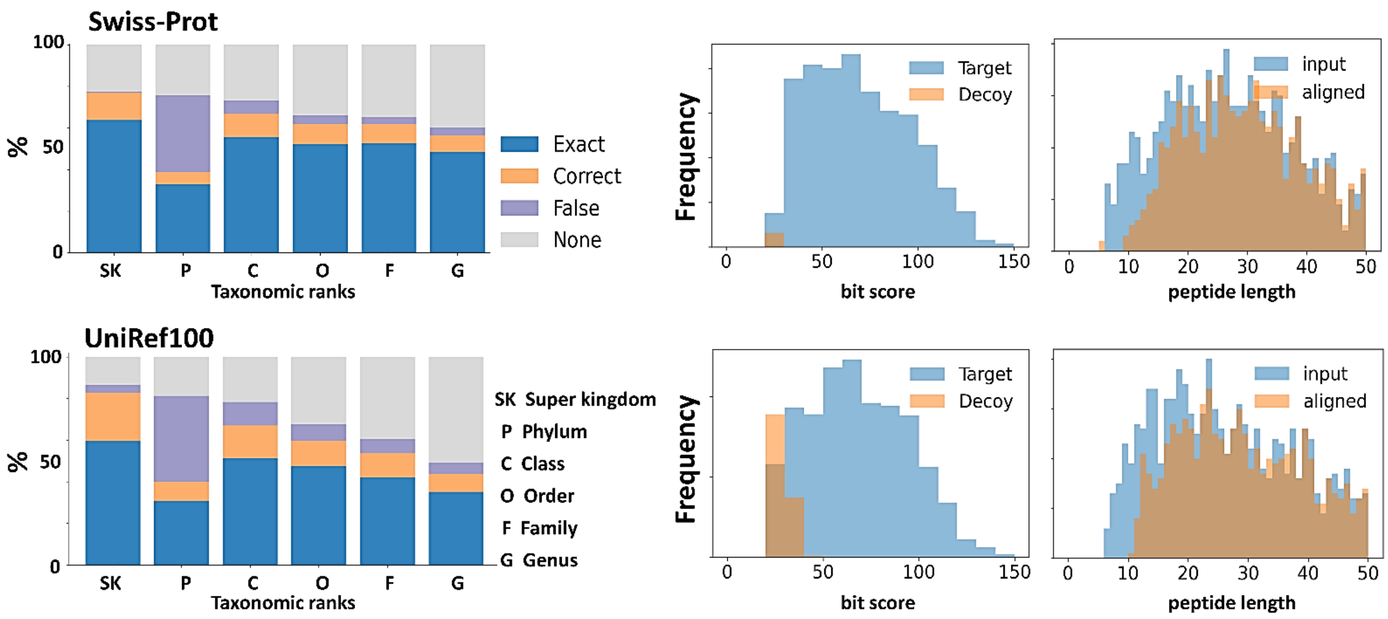


**SI Figure 12.** Sequence alignment recall, precision, random and false matches, bit score and peptide length distribution for combined de novo error dataset #3 (combined_r3.fasta) which covers all types of de novo errors (SI Table 1) with a composition frequency according to previously observed error rates by Muth et al., 2018. Sequences were aligned to the Swiss-Prot and UnRef100 databases.

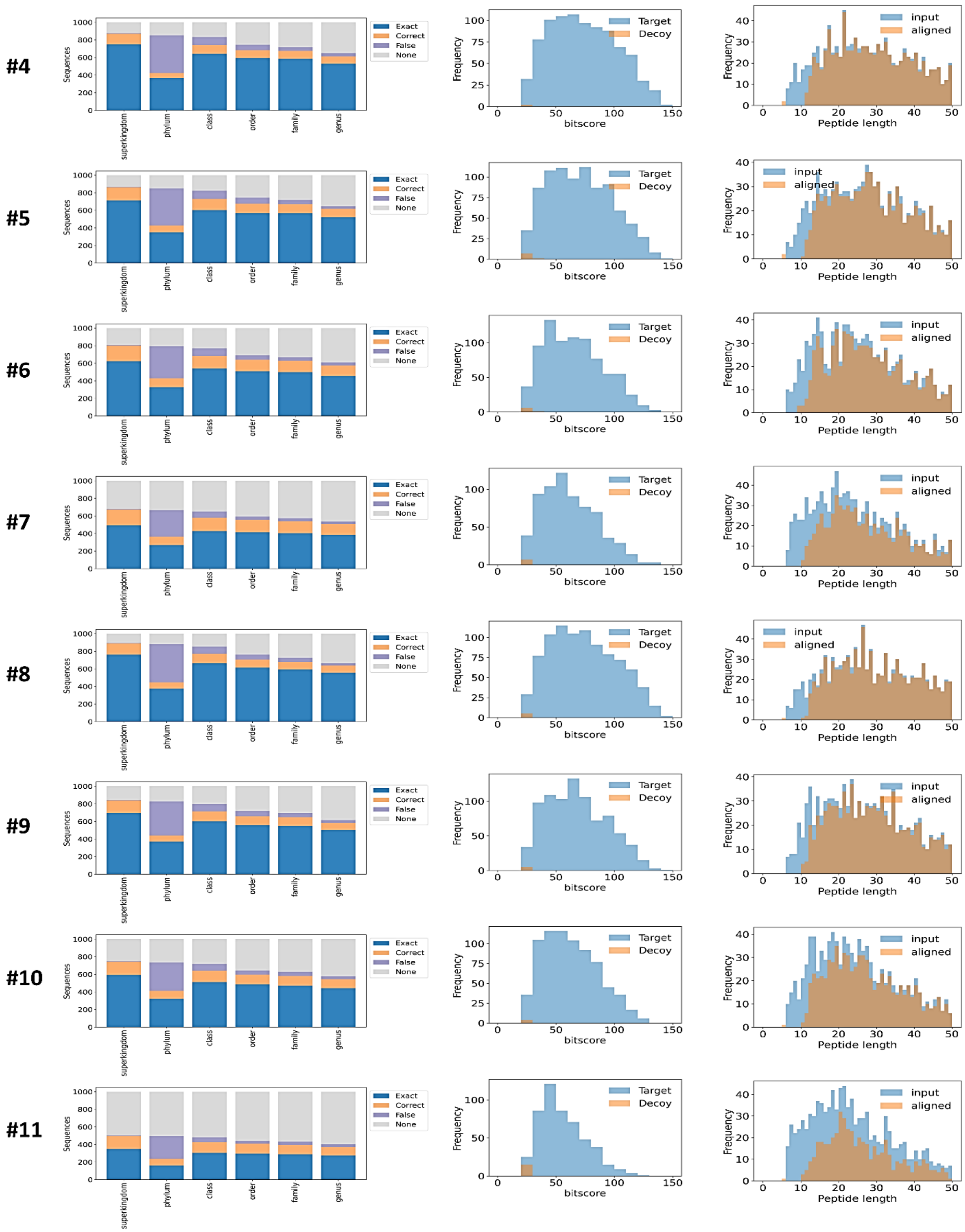


**SI Figure 13.** Sequence alignment performance for datasets #4 = invert_w2c10r1, #5 = invert_w2c1r1, #6 = invert_w2c20r1, #7 = invert_w2c5r1, #8 = invert_w3c10r1, #9 = nvert_w3c1r1, #10 = invert_w3c20r1 and #11 = invert_w3c5r1. Sequences were aligned to the Swiss-Prot database.

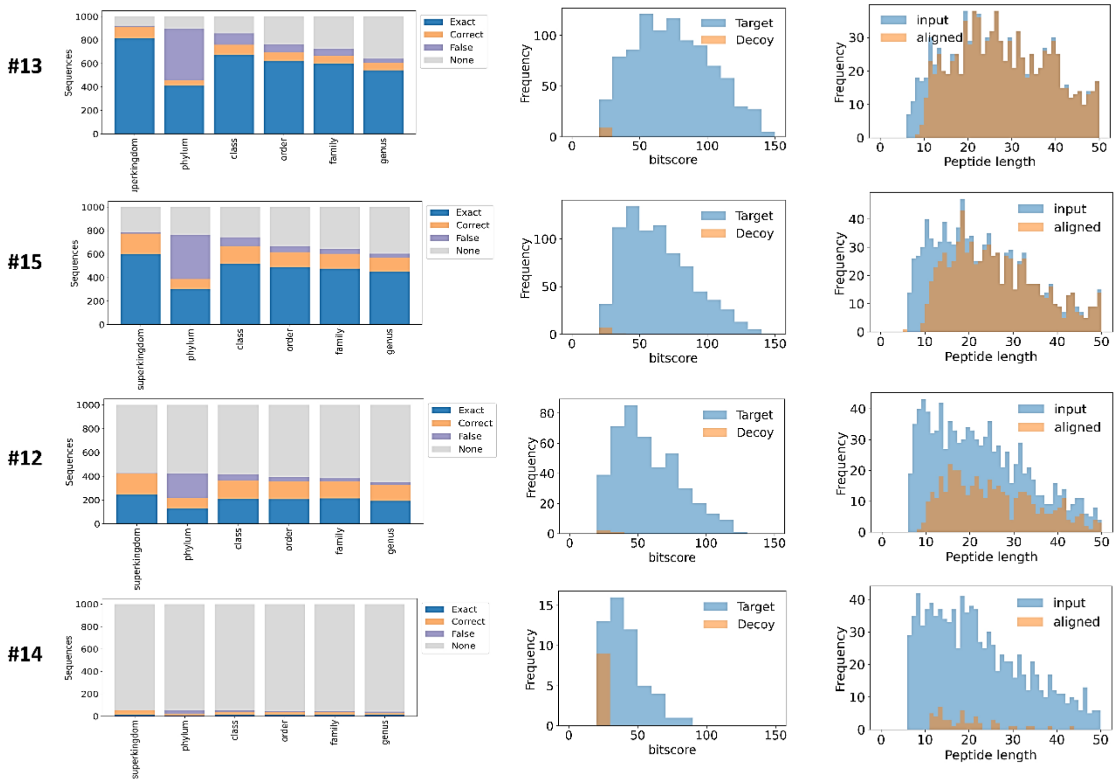


**SI Figure 14.** Sequence alignment performance for datasets #12 = mutate_c10r1, #13 = mutate_c1r1, #14 = mutate_c20r1, #15 = mutate_c5r1. Sequences were aligned to the Swiss-Prot database.

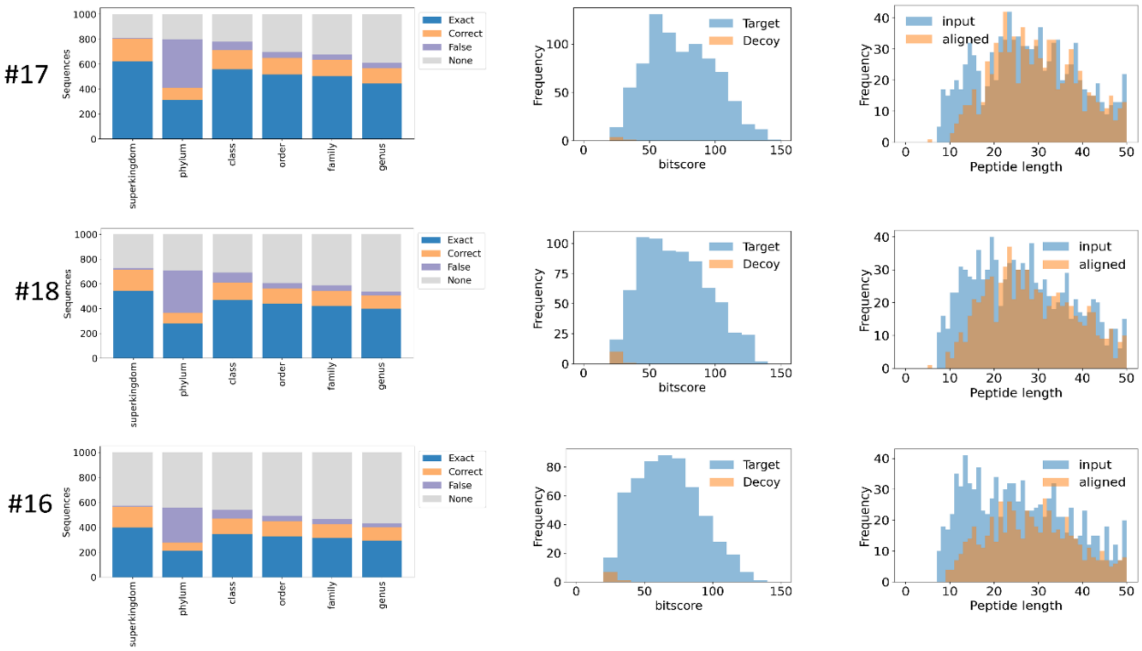


**SI Figure 15.** Sequence alignment performance for datasets #17 = subsitute_s1_2c25r1, #18 = subsitute_s1_2c50r1, #16 = subsitute_s1_2c100r1. Sequences were aligned to the Swiss-Prot database.

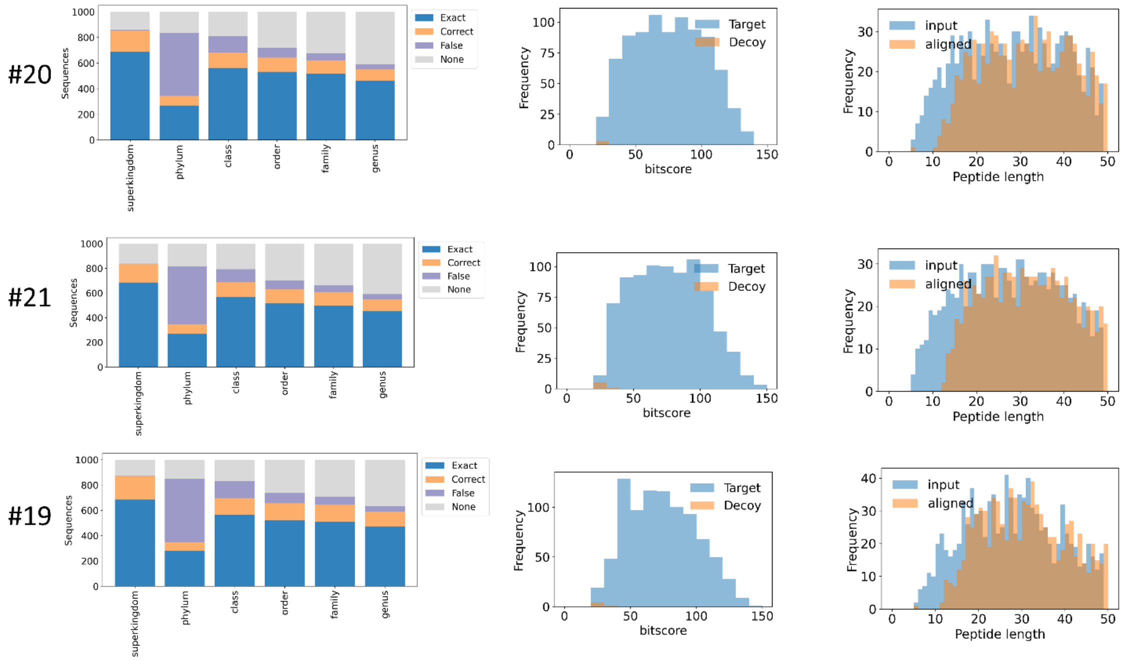


**SI Figure 16.** Sequence alignment performance for datasets #20 = subsitute_s2_1c25r1, #21 = subsitute_s2_1c50r1 and #19 = subsitute_s2_1c100r1. Sequences were aligned to the Swiss-Prot database.

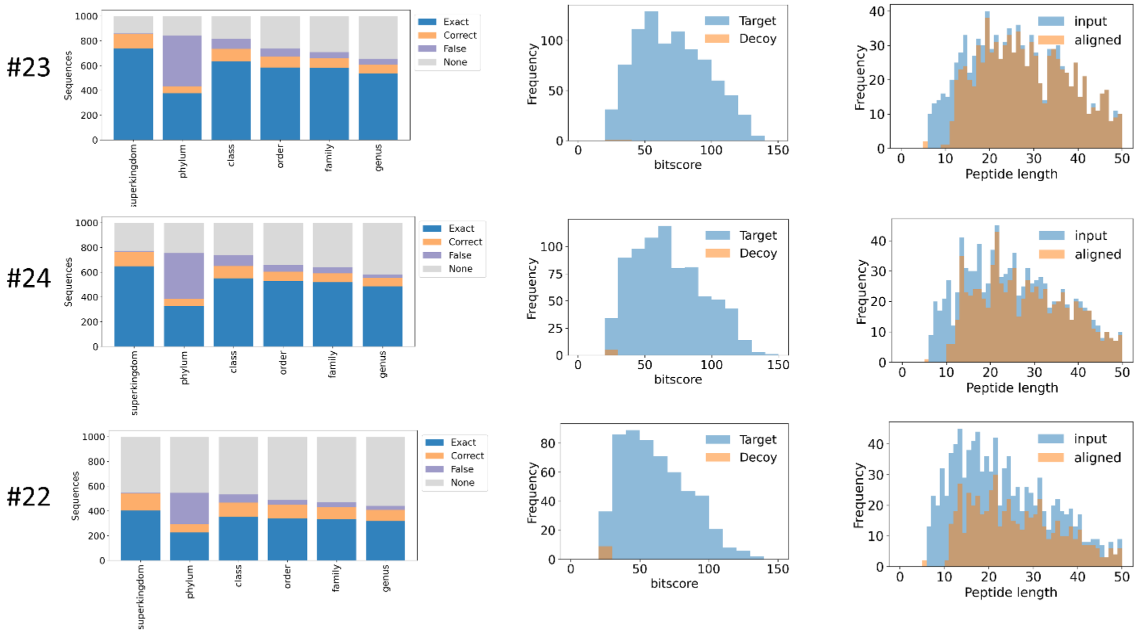


**SI Figure 17.** Sequence alignment performance for dataset #23 = subsitute_s2_2c25r1, #24 = subsitute_s2_2c50r1, #22 = subsitute_s2_2c1SI

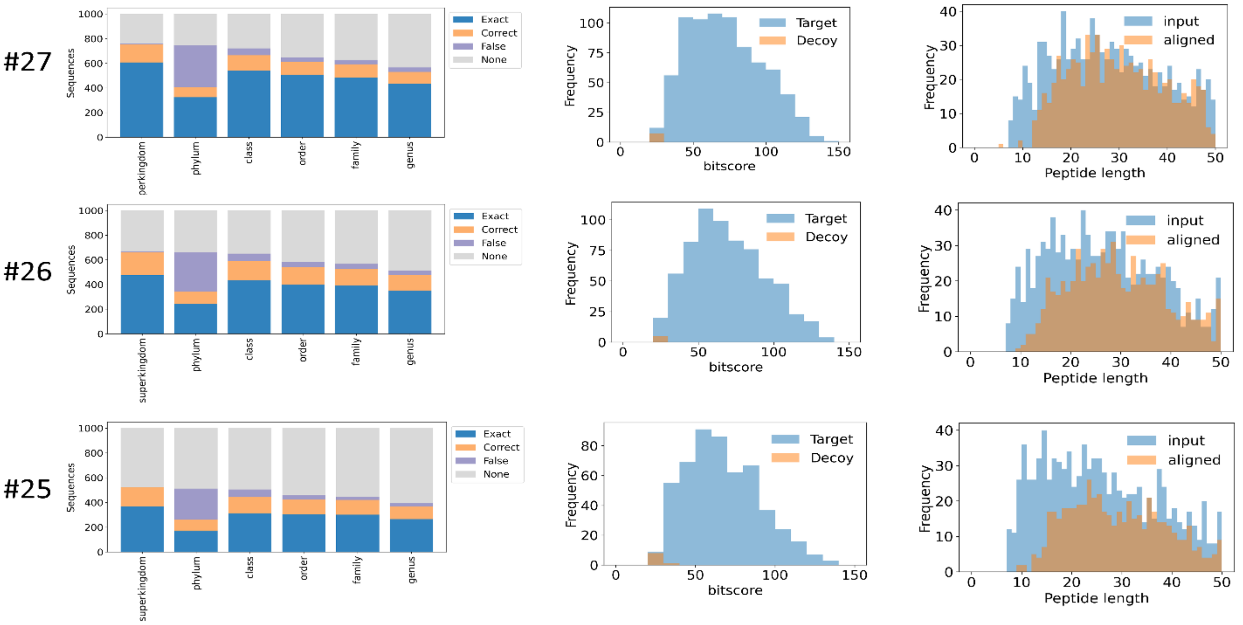


**Figure 18.** Sequence alignment performance for datasets #27 = subsitute_s2_3c25r1, #26 = subsitute_s2_3c50r1, #25 = subsitute_s2_3c100r1. Sequences were aligned to the Swiss-Prot database. Sequences were aligned to the Swiss-Prot database.

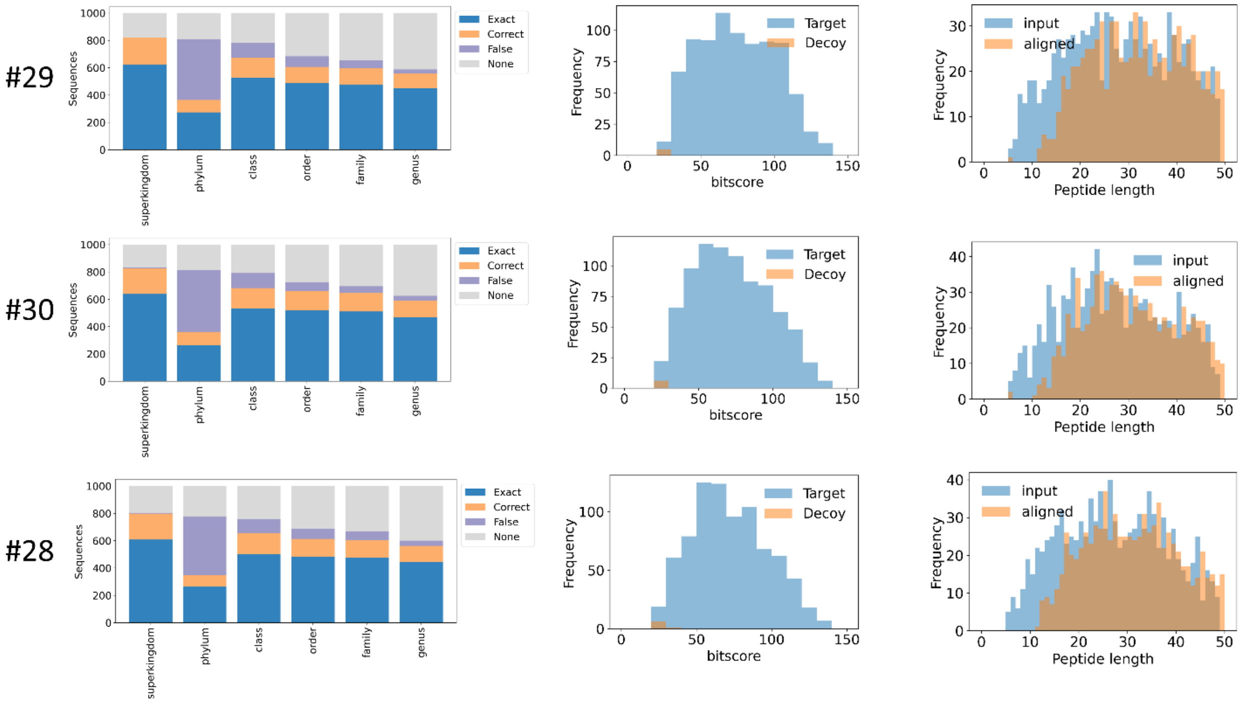


**SI Figure 19.** Sequence alignment performance for datasets #29 = subsitute_s3_2c25r1 #30 = subsitute_s3_2c50r1, #28 = subsitute_s3_2c100r1. Sequences were aligned to the Swiss-Prot database.

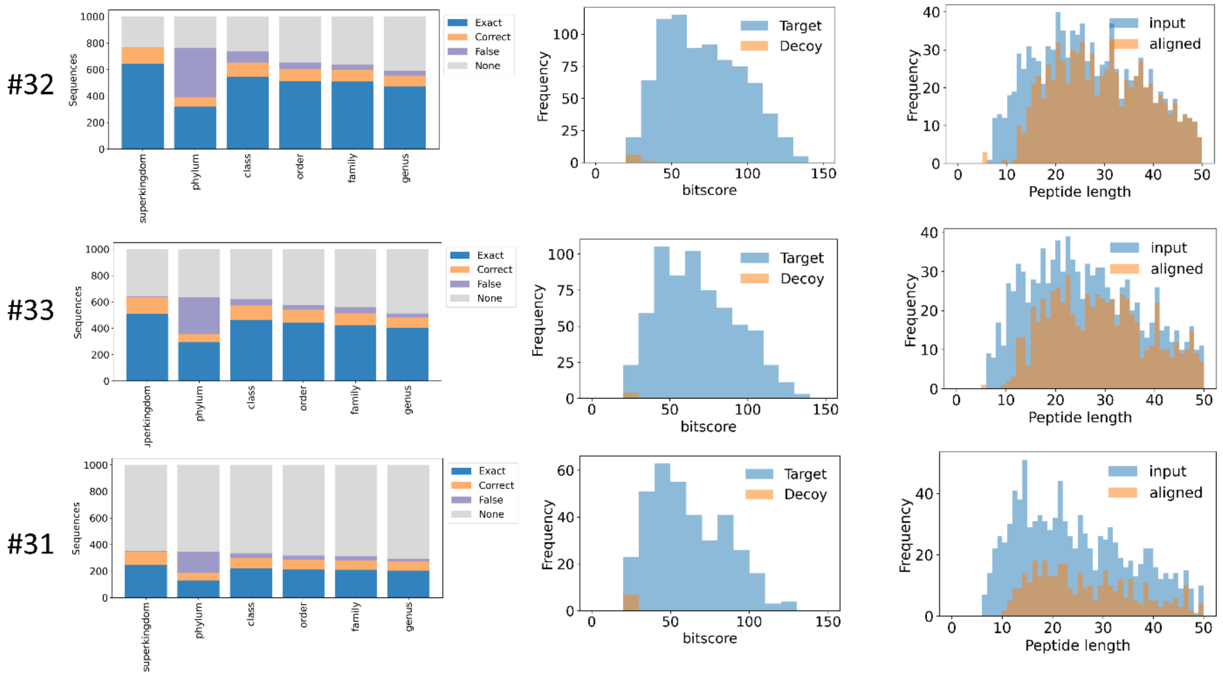


**SI Figure 20.** Sequence alignment performance for datasets #32 = subsitute_s3_3c25r1, #33 = subsitute_s3_3c50r1, #31 = subsitute_s3_3c100r1. Sequences were aligned to the Swiss-Prot database.

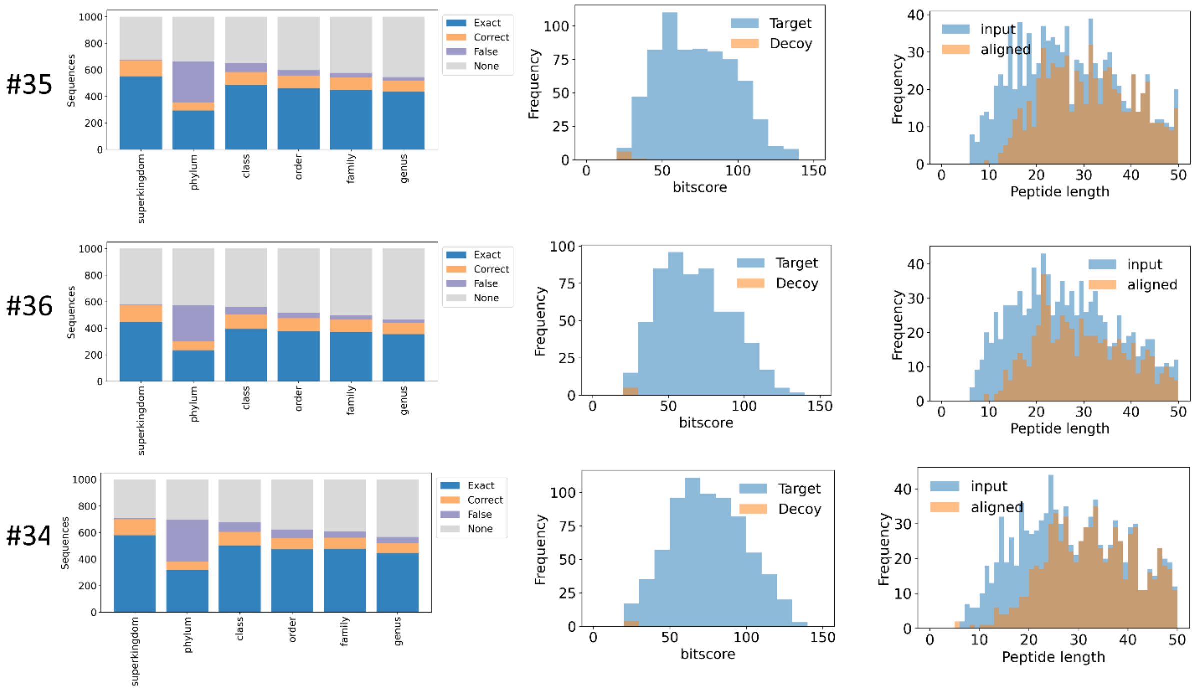


**SI Figure 21.** Sequence alignment performance for datasets #35 = subsitute_s4_4c25r1, #36 = subsitute_s4_4c50r1, #34 = subsitute_s4_4c100r1. Sequences were aligned to the Swiss-Prot database.

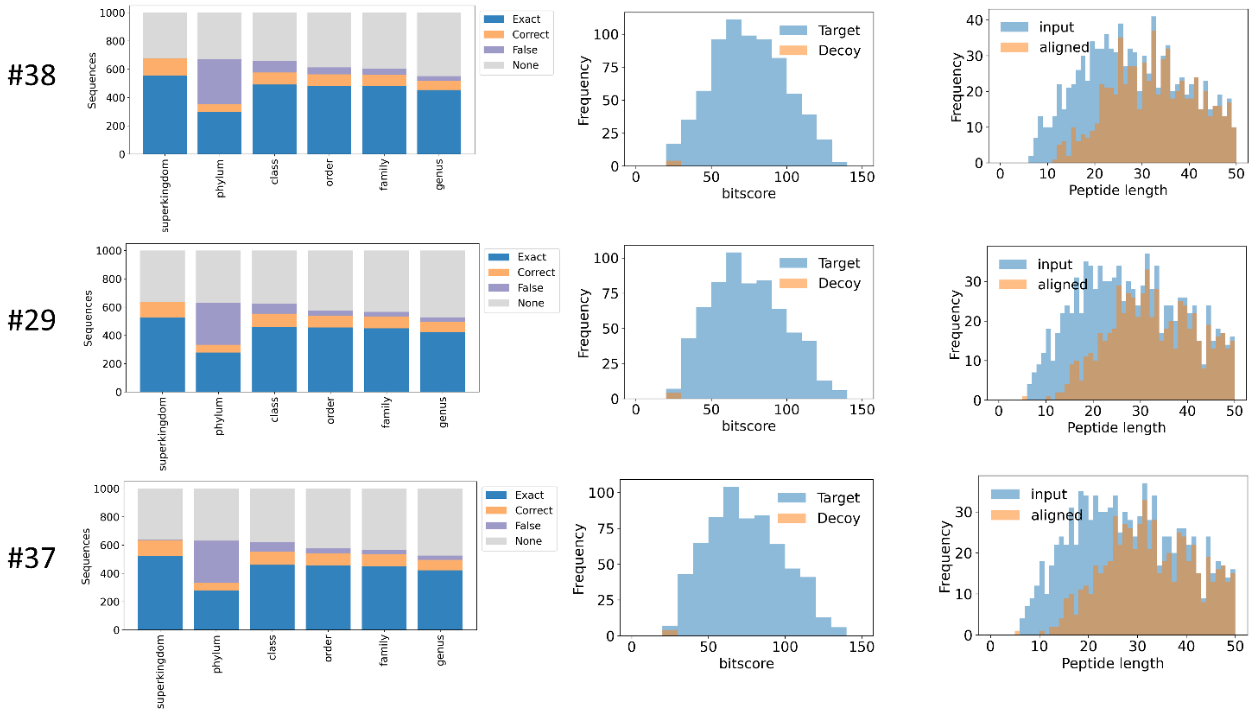


**SI Figure 22.** Sequence alignment performance for datasets #38 = subsitute_s5_5c25r1, #39 = subsitute_s5_5c50r1, #37 = subsitute_s5_5c100r1. Sequences were aligned to the Swiss-Prot database.

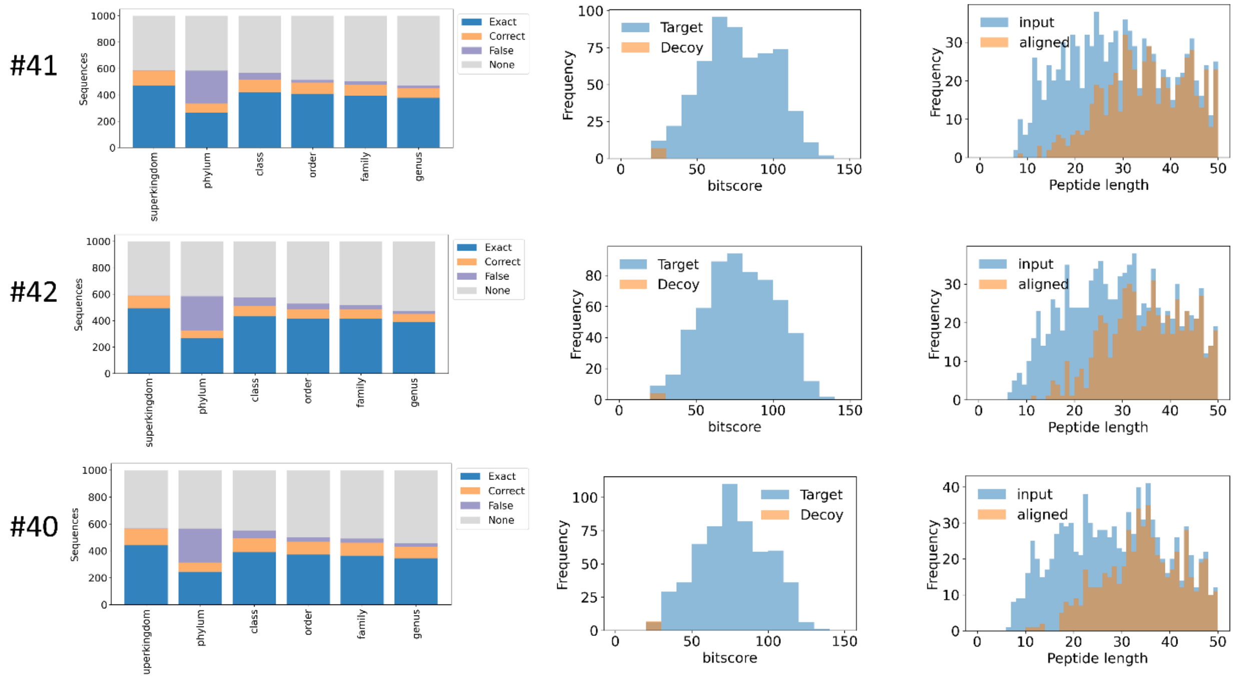


**SI Figure 23.** Sequence alignment performance for datasets #41 = subsitute_s6_6c25r1, #42 = subsitute_s6_6c50r1, #40 = subsitute_s6_6c100r1. Sequences were aligned to the Swiss-Prot database.

### D. Evaluation of NovoLign pipeline for quantitative performance, coverage and % false positives

Following optimization of the DIAMOND alignment procedure for short and minor de novo sequencing errors we aimed to evaluate the NovoLign pipeline using an extensive set of real proteomics data from pure reference strains, synthetic communities, enrichments and complex microbial communities.

In order to evaluate whether the pipeline provides a genuine representation of the community composition we employed 2 synthetic communities with defined taxonomic composition. The first dataset was the “equal protein community” established by Kleiner et al., 2017 (ProteomeXchange server project code PXD006118)[5] and the second synthetic community was the “SIHUMIx” dataset published in the inter laboratory study by Van Den Bossche et al., 2021 (ProteomeXchange server project code PXD023217)[6]. The SIHUMIx is a simplified human intestinal microbiota community which was established by Krause et al., for in vitro studies (Gut Microbes journal, 2020).[7] Furthermore, we used these datasets to optimise the NovoLign pipeline to maximize the number of sequence annotations (“coverage”) by minimizing random as well as false positive annotations. In order to evaluate the available parameters which could impact the final compositional output as well as coverage and false annotations we systematically investigated >80 parameter combinations.

The first parameter we investigated was the de novo quality score for sequences which are subjected to sequence alignment. For PEAKS de novo we investigated ALC (average local confidence) score cutoff levels 50, 70 and 90. Next, we investigated the bit score which is obtained for the individual sequence alignments. There, we investigated different thresholds (20, 25 and 30) for sequences which were further processed, e.g. to obtain a taxonomic composition. Furthermore, most sequences provide high scoring alignments to different taxonomies. Therefore, we investigated also 3 different lowest common ancestor (LCA) algorithms to establish a consensus lineage. The first LCA approach is a “conventional” LCA (“con-LCA”), which strictly aligned the lineages of all sequences above the bit score cutoff to determine the lowest common ancestor.[8] The second LCA approach (“bit-LCA”) is based on the approach used in the BAT tool, which was published by Von Meijenfeldt et al.[9] This approach determines the consensus taxonomy stepwise, where for every taxonomic rank the taxonomic identifier is chosen which accounts for the majority of the total bit score. The third approach is a weighted LCA approach (“WLCA”), which was introduced by Buchfink et al. for metagenomic applications, in 2015.[10] This approach assigns weights to all taxonomies based on their frequency in the alignment results. The consensus lineage for each aligned sequence is determined from taxonomies which combined account for at least 80% of the sum of weights to which the query sequence provided alignments. Finally, NovoLign performs a taxonomic grouping step, where all taxonomic identifiers are grouped to provide a community composition. The sequence counts for each taxonomic group provides then a crude estimator for the abundance of a taxonomy within the community. Nevertheless, in order to avoid reporting extensive lists of taxonomies, with only few sequence counts, we implemented a minimum frequency criterium for the taxonomic grouping (and reporting) step. Thereby, we evaluated thresholds of 5, 10 and 15 sequence counts per taxonomic identifier. Finally, the evaluated parameter combinations are summarized in SI Table S2. All alignments were performed using the UniRef100 reference sequence database (release July 2023; 356800925 sequences; https://ftp.uniprot.org/pub/databases/uniprot/uniref/uniref100/).

**SI Table 2.** Table summarizing the parameters to evaluate the taxonomic profiling accuracy, coverage and % random and false annotations. The image below shows a screenshot of the python code used to loop through the parameters.

| Parameter | Description | Investigated values | | |
|---|---|---|---|---|
| **ALC (%)** | De novo sequence quality/confidence score | 50 | 70 | 90 |
| **bit score** | Statistical significance score for alignment | 20 | 25 | 30 |
| **LCA approach** | Algorithm to determine consensus lineage | con-LCA | bit-LCA | WLCA |
| **Freq cutoff** | Frequency threshold for taxonomic reporting | 5 | 10 | 15 |

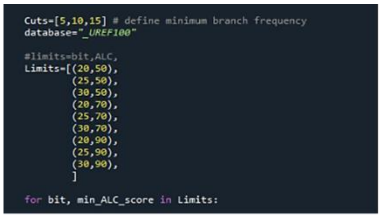

```
Cuts=[5,10,15] # define minimum branch frequency
database="_UREF100"

#limits=bit,ALC,
Limits=[(20,50),
        (25,50),
        (30,50),
        (20,70),
        (25,70),
        (30,70),
        (20,90),
        (25,90),
        (30,90),
        ]

for bit, min_ALC_score in Limits:
```

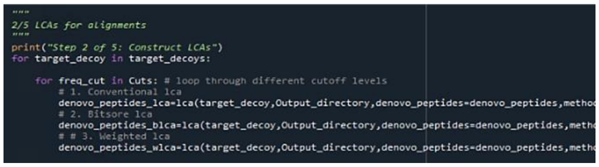

```
"""
2/5 LCAs for alignments
"""
print("Step 2 of 5: Construct LCAs")
for target_decoy in target_decoys:

    for freq_cut in Cuts: # loop through different cutoff levels
        # 1. Conventional lca
        denovo_peptides_lca=lca(target_decoy,Output_directory,denovo_peptides=denovo_peptides,metho
        # 2. Bitsore lca
        denovo_peptides_blca=lca(target_decoy,Output_directory,denovo_peptides=denovo_peptides,meth
        # # 3. Weighted lca
        denovo_peptides_wlca=lca(target_decoy,Output_directory,denovo_peptides=denovo_peptides,meth
```

### D1. Overview of sequence counts and abundance correlation for different parameter combinations

**“Kleiner equal protein”**

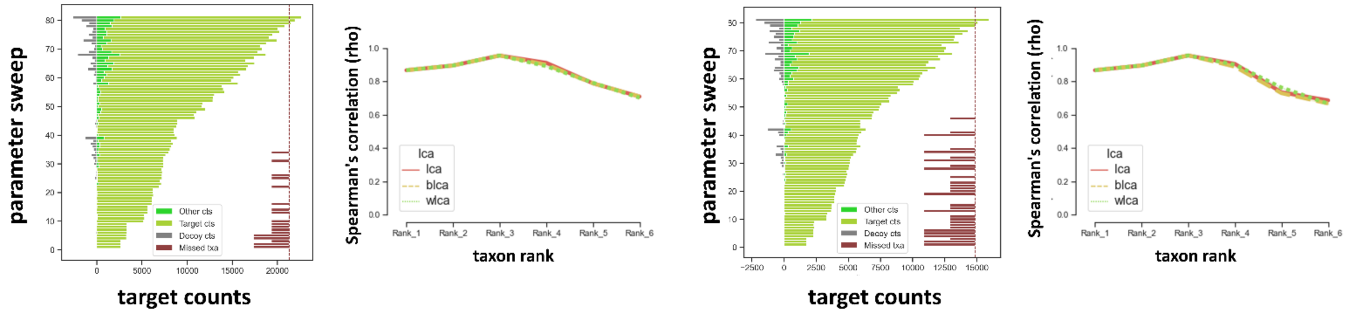


**SIHUMIx**

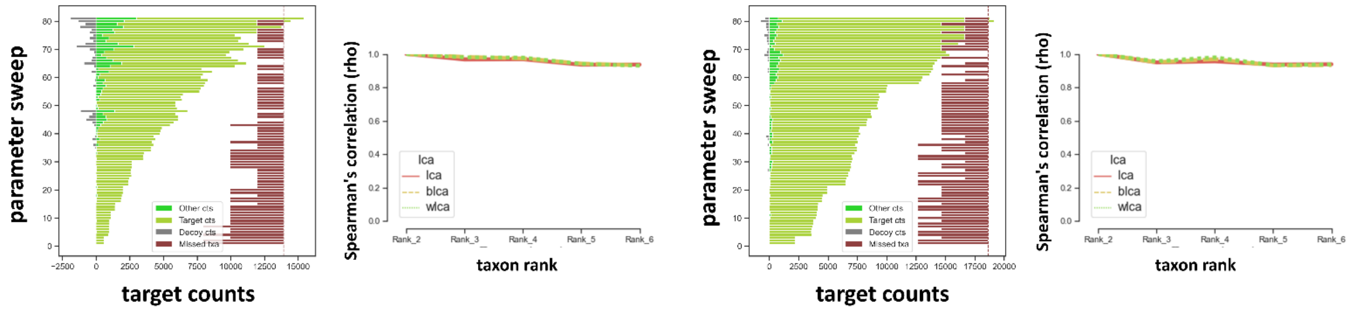


**SI Figure 24 (extension to manuscript Figure 2A).** The bar graphs display the sequence annotations obtained at the family level for the different parameter combinations (see SI Table 1) for both duplicates of the Kleiner equal protein and SIHUMIx communities. The upper pair of graphs details the results for the synthetic equal protein "Kleiner community": Run2_P1_2000ng.raw (left) and Run2_P2_2000ng.raw (right). The lower pair of graphs present the results for the synthetic SIHUMIx samples: S05.raw (left) and S08.raw (right). Each condition is shown as a bar, proportional in size to the number of counts on the x-axis, for expected taxonomies (light green), unexpected other taxonomies (green), and random matches (dark green). The red bars located on the right depict the number of missed taxonomies per condition, with the shortest bar demonstrating a single taxonomy miss. The line graphs show the Spearman's correlation, comparing the abundance of individual taxonomies obtained from the sequence alignment to their expected abundance. Generally, the proportion of decoy (random) and other taxonomies are very low (around <10% for any condition). Missed taxonomies for the Kleiner community were observed only in strict conditions, with very high ALC, bit score, and cutoff thresholds. For the SIHUMIx sample, only taxonomies with an abundance <1% were missed in the majority of conditions. The Spearman correlation graphs demonstrate an excellent correlation between the observed sequence counts abundance and the expected abundance, for both the Kleiner community as well as the SIHUMIx samples. For SIHUMIx, the composition obtained from the database search data was used as a reference. Rank_1 = Superkingdom, Rank_2 = Phylum, Rank_3 = Class, Rank_4 = Order, Rank 5 = Family, Rank_6 = Genus.

### D2. Impact of ALC, bit and cutoff thresholds on % decoy and % other matches

**“Kleiner equal protein”**

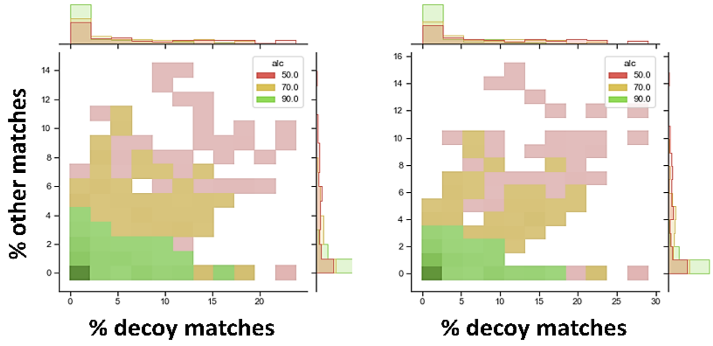


**SIHUMIx**

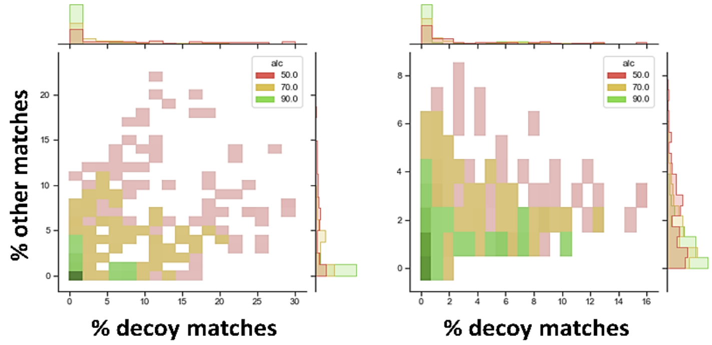


**SI Figure 25:** The graphs show the influence of different ALC filters (50: red, 70: orange, 90: green) on the obtained % other (unexpected taxon) matches (y-axis) and % decoy matches (x-axis). Sequences which are above or equal the ALC threshold are considered for sequence alignment. The upper two graphs depict the outcome for the synthetic equal protein "Kleiner community": Run2_P1_2000ng.raw (left) and Run2_P2_2000ng.raw (right). The lower two graphs show the result for the synthetic SIHUMIx samples: S05.raw (left) and S08.raw (right). Overall, the evaluation reveals a correlation between ALC score and % other matches and % decoy matches. Particularly, % other matches drop to around 10% and 5% when sequences with de novo ALC scores >70 and >90, respectively, are processed.

**“Kleiner equal protein”**

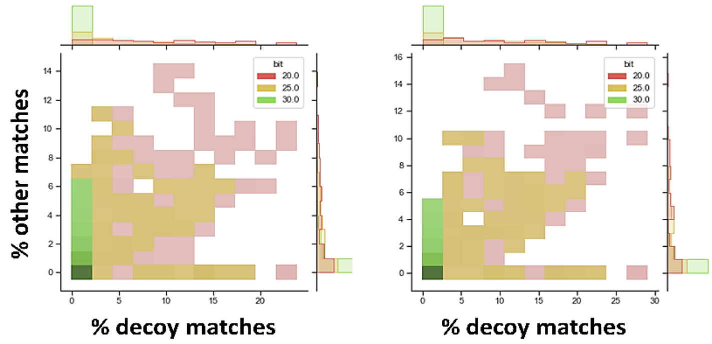


**SIHUMIx**

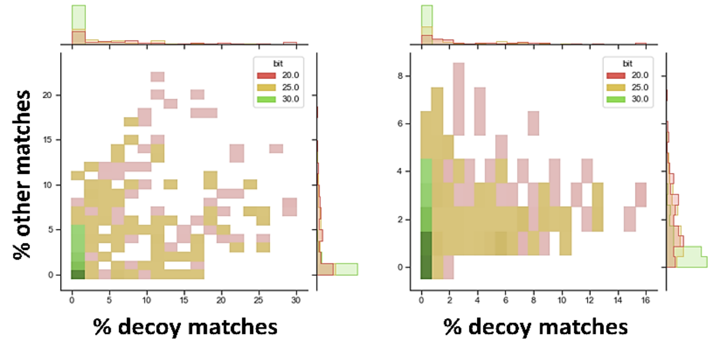


**SI Figure 26:** The graphs show the influence of different alignment bit score filters (20: red, 25: orange, 30: green) on the obtained % other (unexpected taxon) matches (y-axis) and % decoy matches (x-axis). Alignments which are above or equal the bit score threshold are used to determine a consensus lineage for every peptide, using an LCA algorithm. The upper two graphs depict the outcome for the synthetic equal protein "Kleiner community": Run2_P1_2000ng.raw (left) and Run2_P2_2000ng.raw (right). The lower two graphs show the result for the synthetic SIHUMIx samples: S05.raw (left) and S08.raw (right). As expected, both the % decoy and % other decrease when using higher bit score thresholds. However, % decoy matches drop to approximately 1% when only processing alignments with a bit score of 30 or larger.

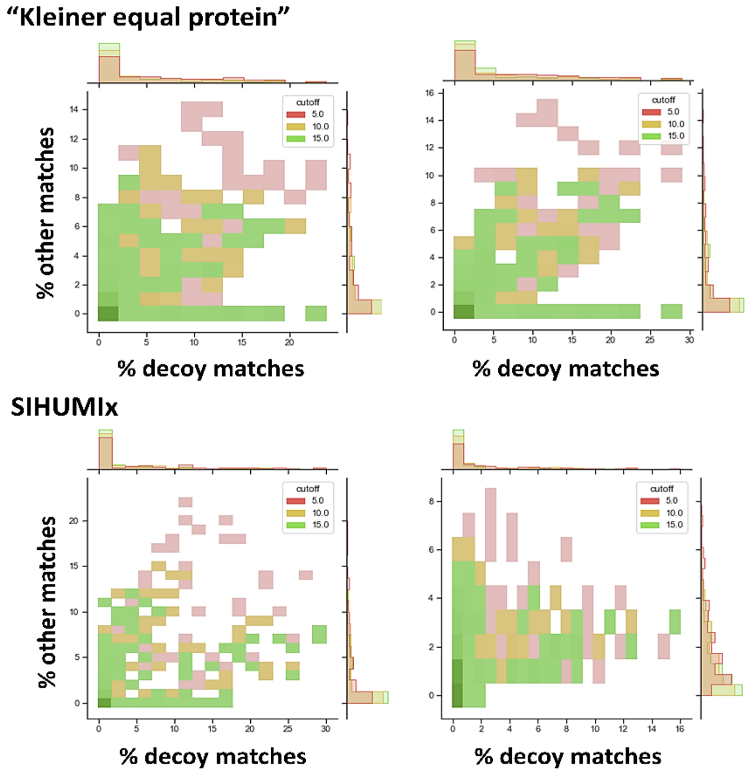


**SI Figure 27:** The graphs show the influence of a taxonomic lineage cutoff threshold (5: red, 10: orange, 15: green), on the %other (unexpected taxon) matches (y-axis) and %decoy matches (x-axis). This cutoff is applied after grouping all taxonomies obtained from the alignments to provide a community composition. The upper two graphs depict the outcome for the synthetic equal protein "Kleiner community": Run2_P1_2000ng.raw (left) and Run2_P2_2000ng.raw (right). The lower two graphs show the result for the synthetic SIHUMIx samples: S05.raw (left) and S08.raw (right). A larger cutoff threshold reduces the %other sequence matches, as expected, since taxonomies only indicated from a few sequence alignments are removed.

### D3. Impact on LCA algorithm on % decoy, % other matches and total taxon counts

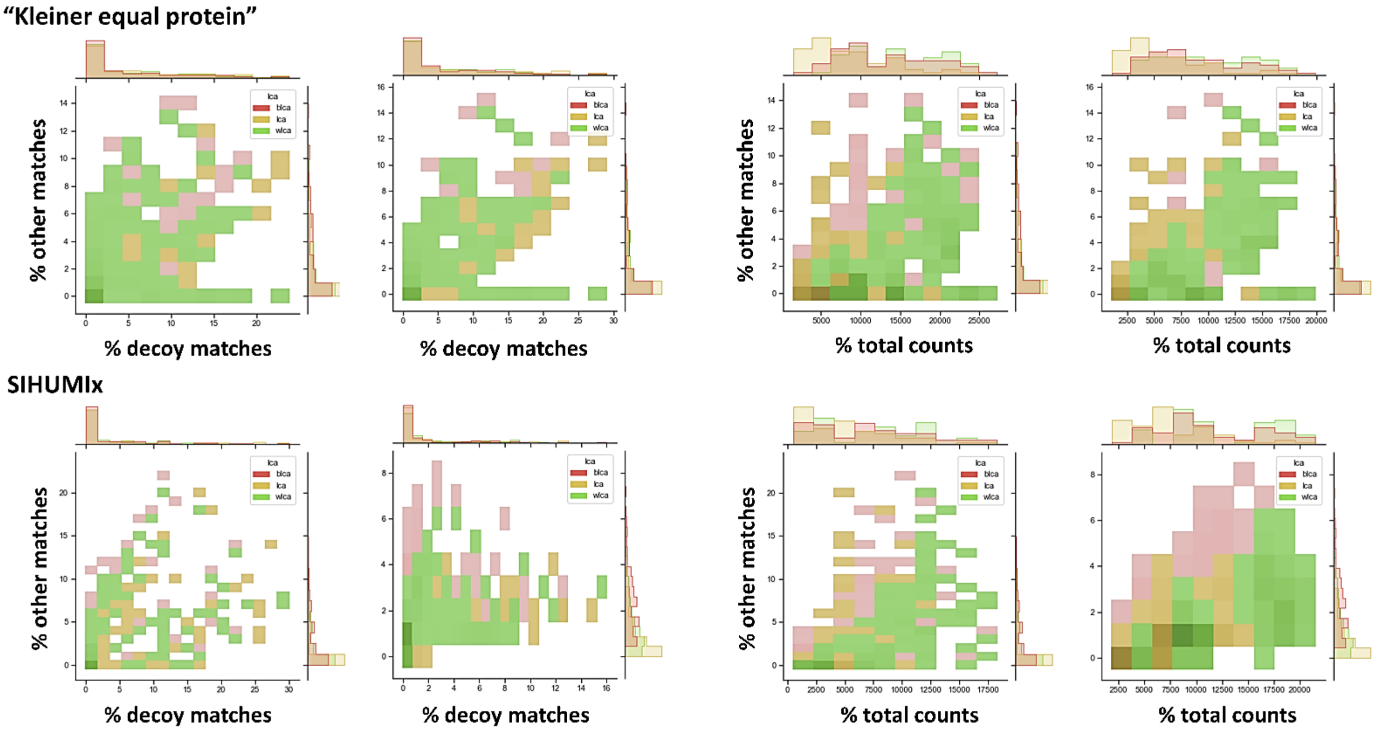


**SI Figure 28:** The graphs show the influence of different LCA algorithms (bit-LCA: red, con-LCA: orange, WLCA: green) on the percentage of other (unexpected) taxa matches (y-axis) and decoy matches (x-axis, two graphs from the left), as well as total counts (x-axis, two graphs from the right). The LCA (lowest common ancestor) algorithm determines a consensus lineage from the TaxIDs of the "subject sequences" that were obtained for each peptide sequence ("query sequence"). The upper four graphs depict the results for the synthetic equal protein "Kleiner community": Run2_P1_2000ng.raw (graphs 1 and 3) and Run2_P2_2000ng.raw (graphs 2 and 4). The lower two graphs show the outcome for the synthetic SIHUMIx samples: S05.raw (graphs 1 and 3) and S08.raw (graphs 2 and 4). The LCA algorithm had no significant impact on the % of other or decoy matches (in the final composition report). Nevertheless, the LCA algorithm had a strong impact on the total number of counts obtained for the observed taxa. The bit-score LCA (bit-LCA) provided a higher number of taxon counts compared to the conventional LCA (con-LCA), while the weighted LCA (WLCA) generally provided more taxon counts than the conventional (con-LCA) and bit-score LCA (bit-LCA).

### D4. Influence of various LCA algorithms and bit scores on total sequence counts

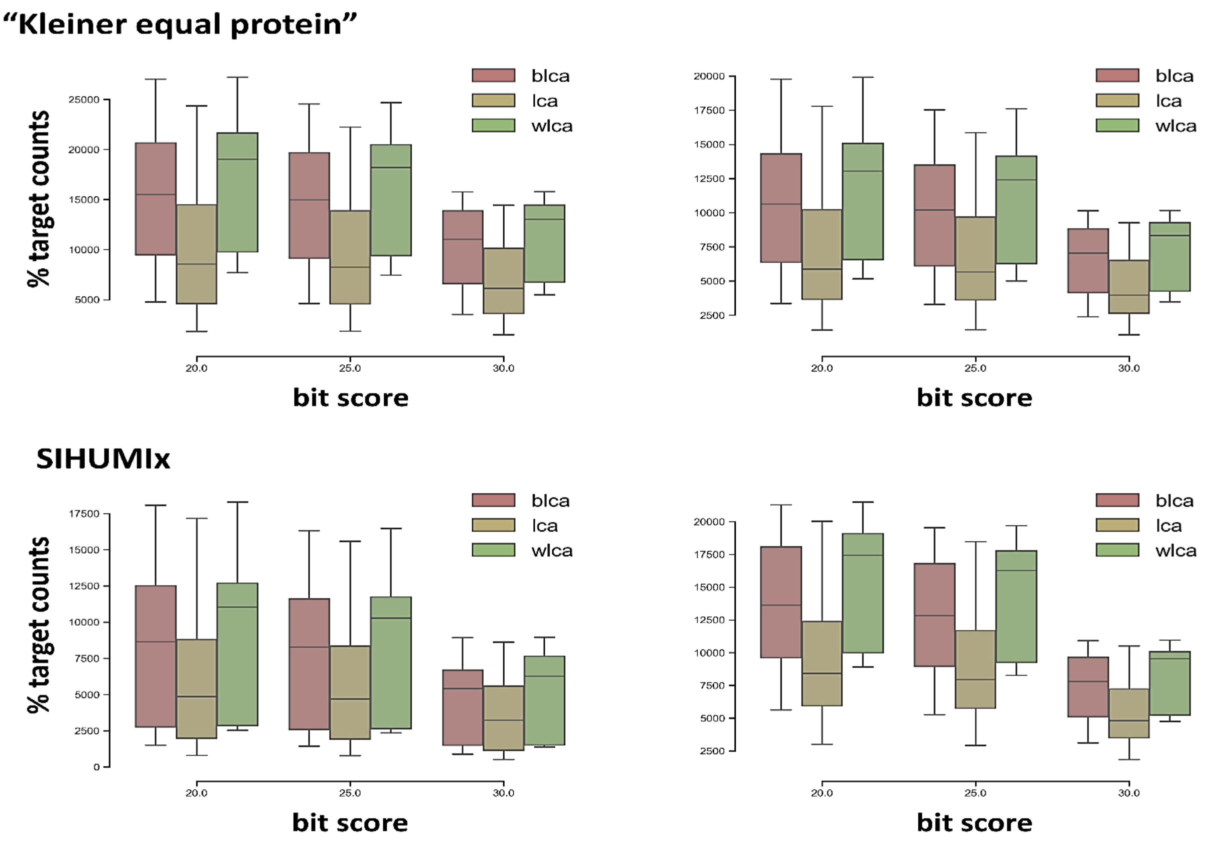


**SI Figure 29:** The graphs show the influence of various Lowest Common Ancestor (LCA) algorithms (bit-LCA: red, con-LCA: orange, WLCA: green) and bit scores on total sequence counts for the observed taxonomies (y-axis). The LCA algorithm determines a consensus lineage from the TaxIDs of the "subject sequences" obtained from each peptide sequence ("query sequence"). The upper graphs depict the results for the synthetic equal protein "Kleiner community": Run2_P1_2000ng.raw (left graph) and Run2_P2_2000ng.raw (right graph). The lower two graphs show the outcomes for the synthetic SIHUMIx samples: S05.raw (left graph) and S08.raw (right graph). Both the LCA algorithm and the bit score significantly affect the total sequence counts for the observed taxonomies. As expected, the higher the bit score threshold, the fewer sequences are assigned to the individual taxonomies. Moreover, bit-score LCA (bit-LCA) yields a higher number of taxon counts in comparison to the conventional LCA (con-LCA), while the weighted LCA (WLCA) generally provides more taxon counts than both the conventional (con-LCA) and bit-score (bit-LCA) LCAs.

### D5. Overview of best-performing NovoLign parameters for the synthetic equal protein "Kleiner community"

**SI Table 3.** Table outlining the best-performing parameters for the synthetic equal protein "Kleiner community" considering a range of qualifiers such as taxonomic coverage, % decoy and other matches <5%, and with fewer than 10 other taxonomic identifications. The weighted LCA (W) and bitscore LCA (BIT) provided more target counts when compared to the corresponding conventional LCA (CON, not shown in table below). Furthermore, while the minimum bitscore threshold of 20 allowed for more target counts, a threshold of 25 reduced the % of unexpected taxa and decoy matches. Additionally, a stricter taxon reporting (frequency) threshold of 15 further excluded less frequent taxonomies. Consequently, in the current study, a minimum ALC of 70 and a bitscore of 25, in conjunction with a taxon reporting threshold of 15 were used. For cases aimed at investigating less abundant microbes, a lower taxon reporting threshold of 5 was selected.

The upper part of the table presents the results from Run2_P1_2000ng.raw, and the lower part of the table shows the results from Run2_P2_2000ng.raw. The complete performance output (for all parameter combinations) can be found in the SI EXCEL Table 2.

| LCA | Cutoff | ALC | bit | 1 | 2 | 3 | 4 | 5 | 6 | 7 | 8 | 9 | 10 | 11 | 12 | 13 | 14 | Other_cts | Target_cts | %other | %decoy |
|---|---|---|---|---|---|---|---|---|---|---|---|---|---|---|---|---|---|---|---|---|---|
| wlca | 15 | 70 | 20 | 3125 | 397 | 729 | 876 | 165 | 794 | 4383 | 305 | 1221 | 2926 | 332 | 1425 | 994 | 690 | 714 | 18362 | 3,743 | 2,48 |
| wlca | 15 | 70 | 25 | 3004 | 379 | 697 | 816 | 156 | 772 | 4194 | 307 | 1174 | 2812 | 305 | 1388 | 965 | 673 | 491 | 17642 | 2,708 | 1,307 |
| blca | 15 | 70 | 20 | 2410 | 350 | 630 | 732 | 84 | 569 | 3587 | 278 | 979 | 2128 | 354 | 1204 | 922 | 624 | 817 | 14851 | 5,214 | 1,647 |
| blca | 15 | 70 | 25 | 2309 | 341 | 613 | 696 | 76 | 564 | 3461 | 281 | 941 | 2059 | 333 | 1175 | 898 | 609 | 547 | 14356 | 3,67 | 1,127 |
| wlca | 15 | 70 | 20 | 1744 | 242 | 565 | 477 | 92 | 449 | 2517 | 344 | 759 | 3493 | 185 | 774 | 514 | 401 | 645 | 12556 | 4,886 | 3,204 |
| wlca | 15 | 70 | 25 | 1681 | 235 | 530 | 436 | 93 | 415 | 2389 | 339 | 712 | 3371 | 166 | 741 | 487 | 391 | 316 | 11986 | 2,569 | 2,138 |
| blca | 15 | 70 | 20 | 1317 | 212 | 495 | 400 | 37 | 333 | 2060 | 312 | 622 | 2655 | 201 | 656 | 490 | 358 | 467 | 10148 | 4,399 | 2,732 |
| blca | 15 | 70 | 25 | 1274 | 211 | 477 | 378 | 36 | 318 | 1969 | 308 | 585 | 2565 | 184 | 634 | 463 | 346 | 317 | 9748 | 3,15 | 1,431 |

| | |
|---|---|
| **1** | **Rhizobiaceae** |
| **2** | **Alteromonadaceae** |
| **3** | **Bacillaceae** |
| **4** | **Burkholderiaceae** |
| **5** | **Chlamydomonadaceae** |
| **6** | **Chromobacteriaceae** |
| **7** | **Enterobacteriaceae** |
| **8** | **Nitrososphaeraceae** |
| **9** | **Paracoccaceae** |
| **10** | **Pseudomonadaceae** |
| **11** | **Roseobacteraceae** |
| **12** | **Staphylococcaceae** |
| **13** | **Xanthomonadaceae** |
| **14** | **Thermaceae** |

### D6. Overview of best-performing NovoLign parameters for the synthetic "SIHUMIx community"

**SI Table 4.** Table outlining the best-performing parameters for the synthetic SIHUMIx community, considering a range of qualifiers such as taxonomic coverage, % decoy and other matches < 5%, and with fewer than 10 other taxonomic identifications. Like for the "Kleiner community", the weighted LCA (W) and bitscore LCA (BIT) provided more target counts compared to the corresponding conventional LCA (CON, not shown). The minimum bitscore threshold of 20 allowed also for more target counts, while a threshold of 25 reduced the % of unexpected taxa and decoy matches. Consequently, a minimum ALC of 70 and a bitscore of 25 combined with a taxon reporting (frequency) threshold of 15 appeared to be the optimal parameters. However, to facilitate the investigation of less abundant microbes a taxon reporting threshold of 5 was more suitable. For example, the members *Bifidobacteriaceae* and *Lactobacillaceae* were only identified when using the lower taxon reporting threshold (not shown in the table). However, based on the database searching results, both are very low abundant, at approximately 0.15 and 0.05%, respectively.

The upper part of the table presents the results from S05.raw, and the lower part of the table shows the results from S08.raw. The complete performance output for all parameter combinations can be found in the SI EXCEL Table 2.

| LCA | Cutoff | ALC | bit | 1 | 2 | 3 | 4 | 5 | 6 | 7 | Other_cts | Target_cts | %other | %decoy |
|---|---|---|---|---|---|---|---|---|---|---|---|---|---|---|
| wlca | 15 | 70 | 20 | 5333 | 1285 | 608 | 2247 | 346 | 0 | 0 | 503 | 9819 | 4,873 | 1,976 |
| wlca | 15 | 70 | 25 | 5082 | 1246 | 532 | 2160 | 261 | 0 | 0 | 385 | 9281 | 3,983 | 1,086 |
| blca | 15 | 70 | 20 | 4253 | 733 | 486 | 2045 | 93 | 0 | 0 | 518 | 7610 | 6,373 | 0,812 |
| blca | 15 | 70 | 25 | 4071 | 724 | 424 | 1979 | 80 | 0 | 0 | 419 | 7278 | 5,444 | 0,689 |
| wlca | 15 | 70 | 20 | 9185 | 1579 | 1373 | 4165 | 392 | 0 | 0 | 337 | 16694 | 1,979 | 1,227 |
| wlca | 15 | 70 | 25 | 8573 | 1474 | 1277 | 3927 | 348 | 0 | 0 | 192 | 15599 | 1,216 | 0,222 |
| blca | 15 | 70 | 20 | 7408 | 958 | 1008 | 3637 | 81 | 0 | 0 | 593 | 13092 | 4,333 | 0,548 |
| blca | 15 | 70 | 25 | 6954 | 893 | 938 | 3471 | 72 | 0 | 0 | 469 | 12328 | 3,665 | 0 |

| | |
|---|---|
| **1** | **Bacteroidaceae** |
| **2** | **Coprobacillaceae** |
| **3** | **Enterobacteriaceae** |
| **4** | **Lachnospiraceae** |
| **5** | **Clostridiaceae** |
| **6** | **Bifidobacteriaceae** |
| **7** | **Lactobacillaceae** |

### D7. Abundance profiles for the synthetic "Kleiner community" for different LCAs and frequency cutoffs

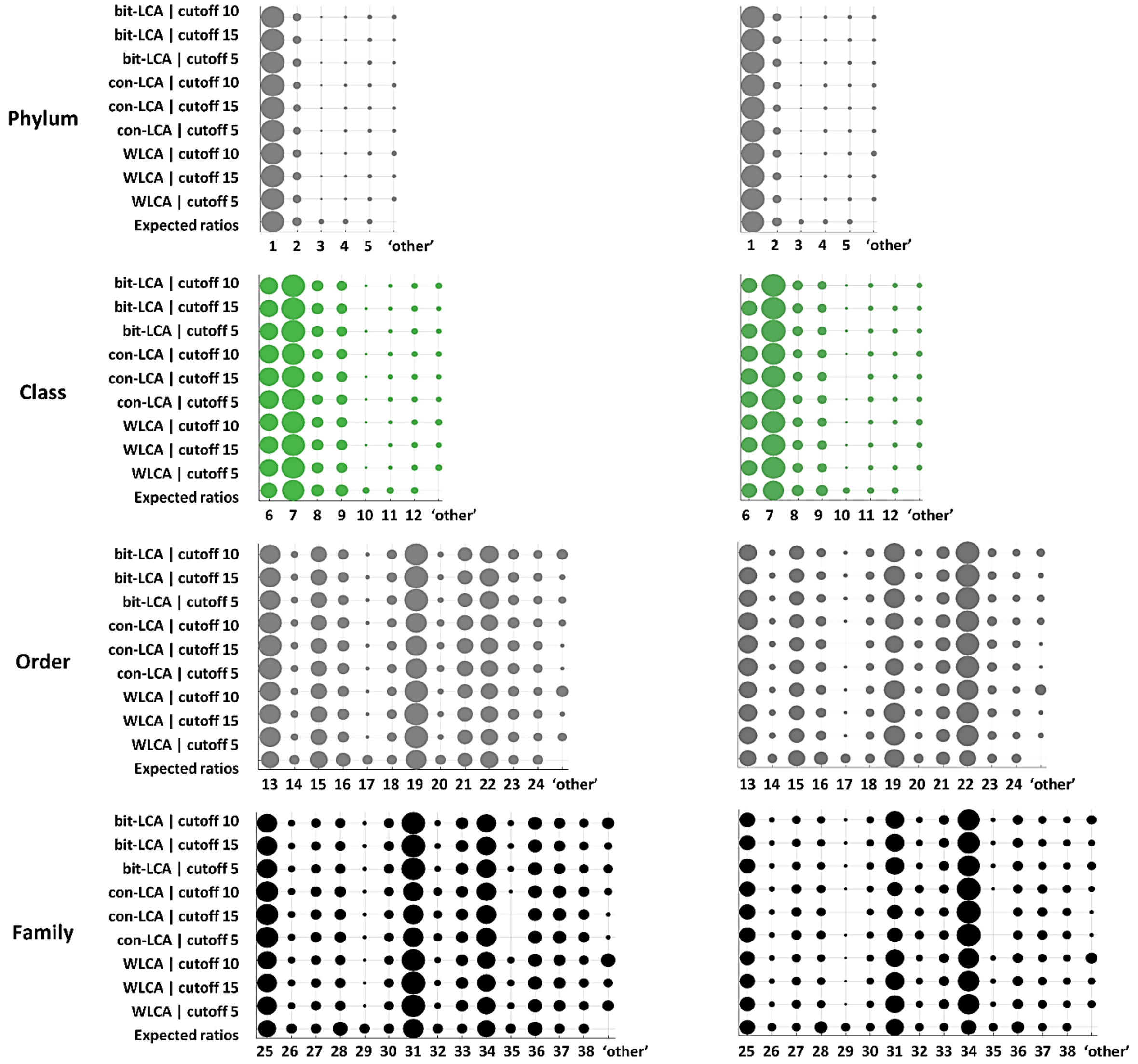


**SI Figure 30:** The circle graphs illustrate the abundance profiles for the synthetic "Kleiner equal protein community" at the different taxonomic ranks of Phylum, Class, Order, and Family, and different LCA algorithms (con-LCA, bit-LCA, and WLCA) and taxonomic frequency cutoff thresholds (5, 10, and 15). The left graphs depict the results for the sample Run2_P1_2000ng.raw (left graph) and the right graph the sample Run2_P2_2000ng.raw. The numbers below the circles label the taxonomic identifiers: 1) Pseudomonadota; 2) Bacillota; 3) Chlorophyta; 4) Nitrososphaerota; 5) Deinococcota; 6) Alphaproteobacteria; 7) Gammaproteobacteria; 8) Bacilli; 9) Betaproteobacteria; 10) Chlorophyceae; 11) Nitrososphaeria; 12) Deinococci; 13) Hyphomicrobiales; 14) Alteromonadales; 15) Bacillales; 16) Burkholderiales; 17) Chlamydomonadales; 18) Neisseriales; 19) Enterobacterales; 20) Nitrososphaerales; 21) Rhodobacterales; 22) Pseudomonadales; 23) Xanthomonadales; 24) Thermales; 25) *Rhizobiaceae*; 26) *Alteromonadaceae*; 27) *Bacillaceae*; 28) *Burkholderiaceae*; 29) *Chlamydomonadaceae*; 30) *Chromobacteriaceae*; 31) *Enterobacteriaceae*; 32) *Nitrososphaeraceae*; 33) *Paracoccaceae*; 34) *Pseudomonadaceae*; 35) *Roseobacteraceae*; 36) *Staphylococcaceae*; 37) *Xanthomonadaceae*; 38) *Thermaceae*. The "Expected ratios" show the expected equal protein abundance ratios.

### D8. Abundance profiles for the synthetic SIHUMIx community for different LCAs and frequency cutoffs

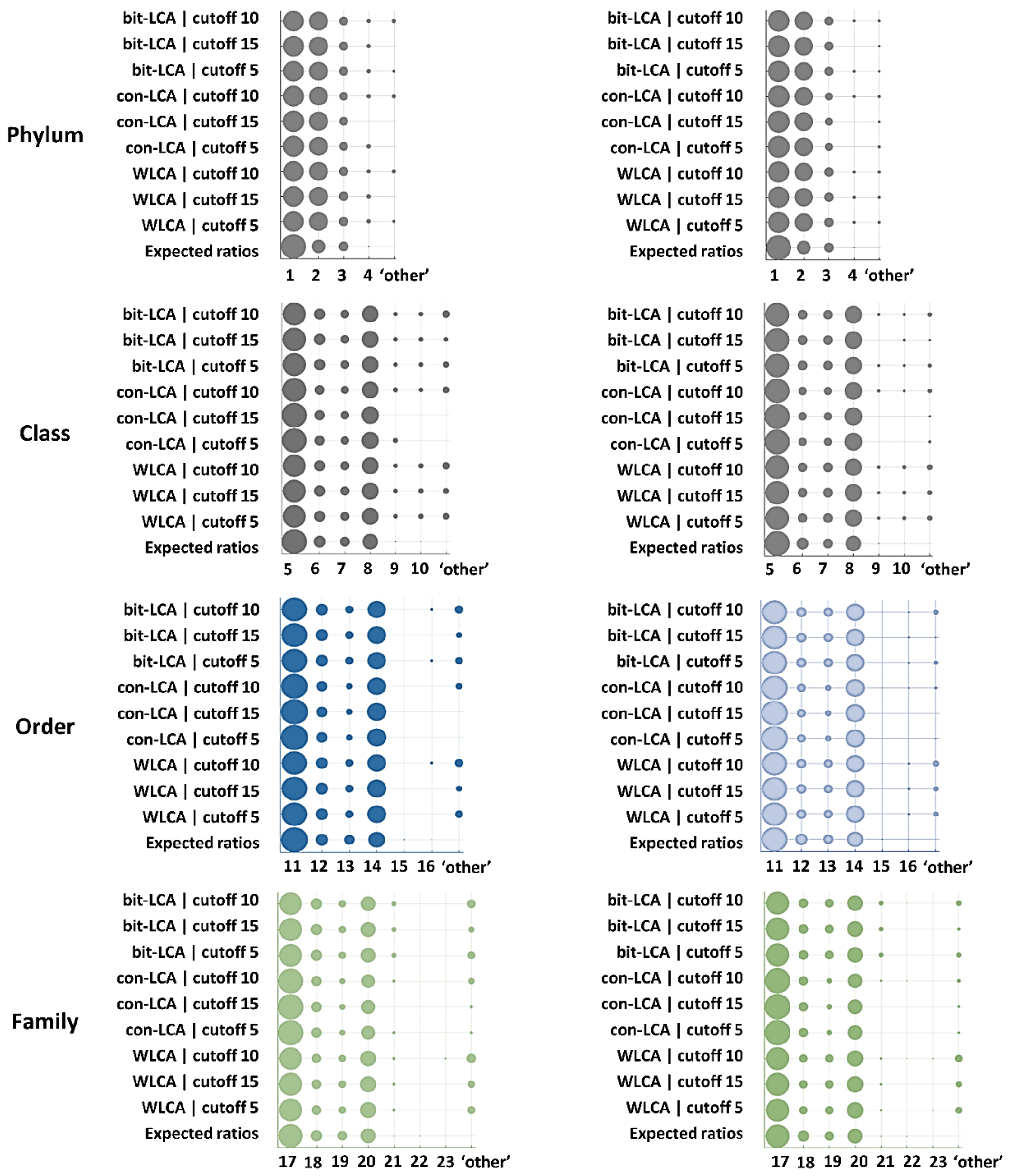


**SI Figure 31:** The circle graphs illustrate the abundance profiles for the synthetic SIHUMIx community at the different taxonomic ranks of Phylum, Class, Order, and Family, and different LCA algorithms (con-LCA, bit-LCA, and WLCA) and taxonomic frequency cutoff thresholds (5, 10, and 15). The left graphs depict the results for the sample S05.raw and the right graphs the sample S08.raw. The numbers below the circles indicate the following taxonomic identifiers: 1) Bacteroidetes; 2) Bacillota; 3) Pseudomonadota; 4) Actinomycetota; 5) Bacteroidia; 6) Erysipelotrichia; 7) Gammaproteobacteria; 8) Clostridia; 9) Actinomycetes; 10) Bacilli; 11) Bacteroidales; 12) Erysipelotrichales; 13) Enterobacterales; 14) Eubacteriales; 15) Bifidobacteriales; 16) Lactobacillales; 17) *Bacteroidaceae*; 18) *Coprobacillaceae*; 19) *Enterobacteriaceae*; 20) *Lachnospiraceae*; 21) *Clostridiaceae*; 22) *Bifidobacteriaceae*; 23) *Lactobacillaceae*. The "Expected ratios" show the expected abundance ratios which was determined by database searching using "DB1" (*Van Den Bossche et al., 2022*).

### D9. Comparison of sequence coverage between NovoLign and NovoBridge

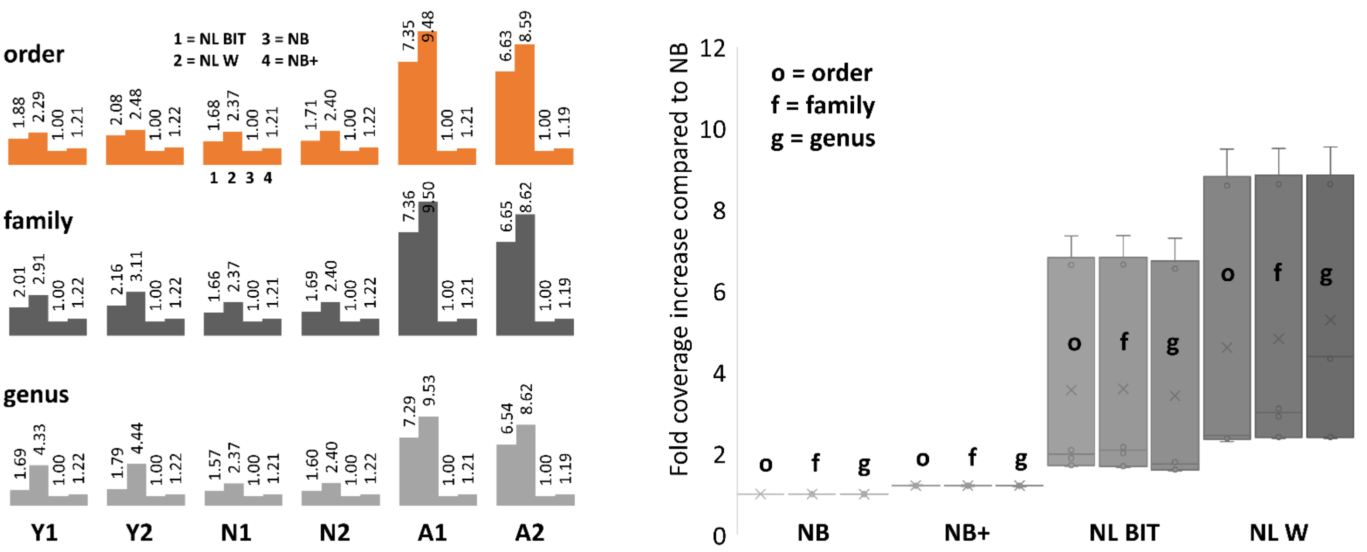


**SI Figure 32.** The left graph illustrates the increase in fold coverage compared to NovoBridge (NB, exact sequence matches only) and NB+ (exact sequence matches, including missed cleavages) for Saccharomyces cerevisiae (Y1 and Y2), Nitrospira moscoviensis (N1 and N2), and Aeromonas bestiarum (A1 and A2). The data processing utilized duplicate datasets (denoted as 1 and 2). The left boxplot displays the average fold increase in peptide sequences annotated with the target species, relative to NB, across different taxonomic levels. On average, NB+ demonstrated an approximate 1.2-fold increase, NL BIT an approximate 3.5-fold increase, and NL W an approximate 5-fold increase. NB = NovoBridge, NB+ = NovoBridge+, NL BIT = NovoLign bit score LCA, and NL W = NovoLign weighted LCA.

### D10. Processing time performance of NovoLign pipeline

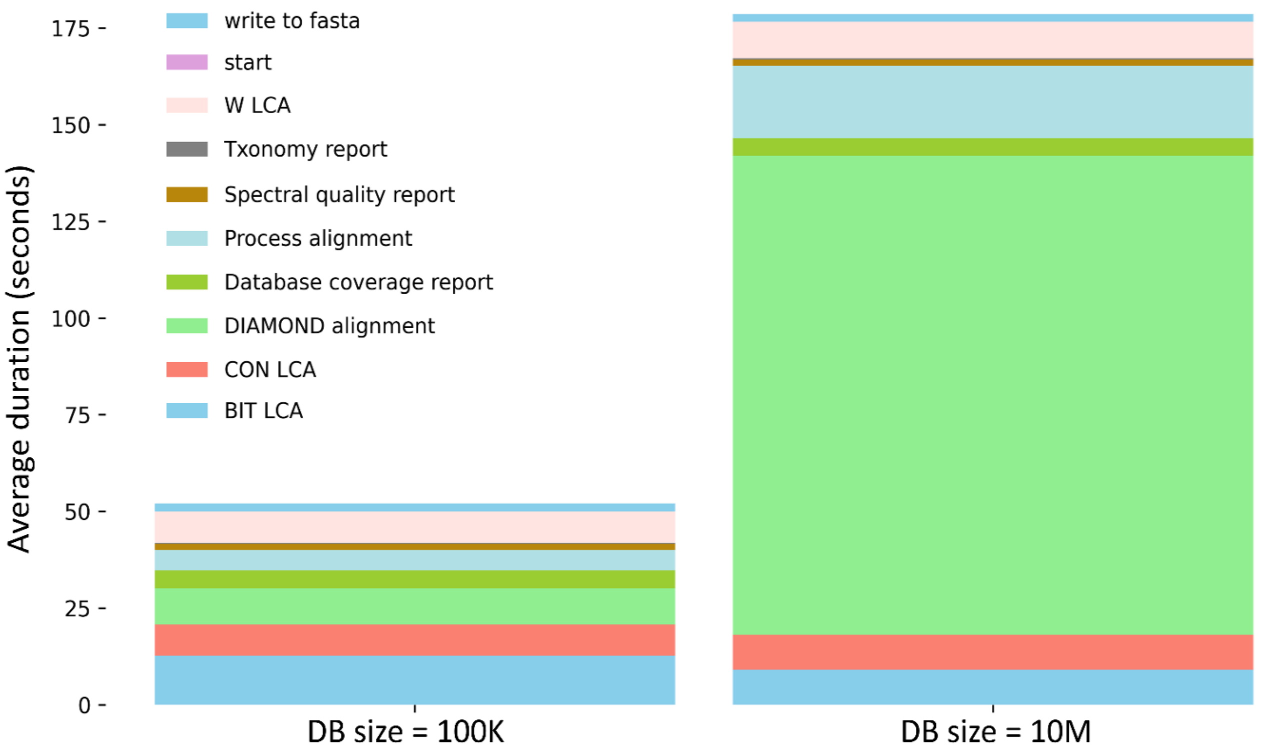


**SI Figure 33:** The processing times for the yeast de novo test dataset through the entire NovoLign pipeline were recorded. This included the initial steps (importing modules, defining parameters, and importing files), writing to FASTA (creating a cleaned de novo sequence dataset, a decoy dataset, and writing these to a FASTA file), DIAMOND alignment (aligning sequences from the FASTA file, including decoys), processing alignment (importing alignments into Python and performing score filtering), and executing taxonomic grouping using different approaches (CON LCA = conventional LCA, BIT LCA = bit score LCA, W LCA = weighted LCA). It also involved generating taxonomy, spectral quality, and database coverage reports. The sequences were aligned against sequence databases containing randomized sequences of 100 thousand (DB size = 100K) and 10 million (DB size = 10M) sequences. Shuffled sequences were created by randomly sampling SwissProt entries and then randomly reshuffling blocks of 10 consecutive amino acids. The yeast test dataset contained 29,184 cleaned de novo sequences. Including decoy sequences, this amounted to 58,368 processed sequences. Performance benchmarking was conducted using the Windows Performance Monitor, tracking the Total % Processor Time on a Precision 3581 Dell workstation with an i7-13700H processor and 32 GB of RAM.

Processing the same yeast de novo dataset with the complete UniRef100 database (containing 352,965,587 sequences) and the complete SwissProt database (containing 567,413 sequences) takes approximately 32.4 and 0.7 minutes, respectively (on a desktop with an Intel(R) Core(TM) i7-7700K and 32 GB RAM).
